# Stress-induced Metabolic Remodeling of Adipose and Brain Tissue Revealed by Positron Emission Tomography

**DOI:** 10.64898/2026.09.09.749803

**Authors:** Maximilian Krisch, Öykü Özer, Jayne Louise Wilson, Sabine Macho-Maschler, Paul Ettel, Hanka Vrban, Alexander Grentner, Barbara Katharina Geist, Clara T.S. Mertel, Usevalad Ustsinau, Oana Kulterer, Joachim Friske, Thomas Wanek, Claudia Kuntner, Thomas Helbich, Stefan Grünert, Rupert Palme, Florian W. Kiefer, Thomas Weichhart, Cécile Philippe, Marcus Hacker, Chrysoula Vraka

## Abstract

Stress impacts our health and triggers physiological adaptations, yet the metabolic programs engaged during stress remain incompletely understood. To fill this knowledge gap, we utilize total-body positron emission tomography (PET), multi-OMICS, and endocrine profiling to assess how various murine stress models affect systemic metabolic remodeling. We find that acute immobilization and surgery activate brown adipose tissue as part of the stress response, independently of hypothermia, thereby acting as a highly stress-sensitive metabolic hub. Additionally, we identify stress-specific hypo- and hypermetabolic signatures in different brain regions, and distinguish networks between brain and adipose tissue depots across the different stress groups. Our work presents a novel perspective on stress and its mobilization of metabolic resources and identifies PET imaging of brain and adipose tissue as a valuable, minimally invasive technique for tracking metabolic stress responses in mice, with relevance for animal welfare and disease models, and translational impact for mental health studies and preventive medicine.

## Introduction

Excessive stress exposure is known to have adverse effects on health, increasing the risks of cardiovascular events^1,2^, metabolic diseases^3^, therapy resistance^4^, and a poorer survival prognosis^5^. In contrast, growing evidence suggests that acute and moderate stress, such as brief cold exposure, can have beneficial effects, including enhanced metabolic function^6^, cardiometabolic health^7^, and reduced tumor progression^8^. These opposing effects highlight the duality of the stress response, emphasizing the need for unbiased, robust, and accurate stress assessment methods that also capture its physiological impact.

Stress elicits adaptive responses to homeostatic challenges via two primary axes. The hypothalamic-pituitary-adrenal (HPA) axis triggers the release of glucocorticoids (GCs), which regulate multiple critical biological processes via glucocorticoid receptors, including cell growth and differentiation^9^, metabolism^10^, cognition^11^, cardiovascular function^12^, and immunity^13,14^. Although prone to inaccurate validation due to the circadian rhythm^15^ and the need for longitudinal sampling^16^, GC levels remain the reference standard for assessing stress^17^. Complementing the HPA axis, the sympathoadrenal medullary axis (SAM) rapidly releases catecholamines, such as epinephrine and norepinephrine, triggering the fight-or-flight response. In peripheral tissues, catecholamines mediate physiological and metabolic responses via α- and β-adrenergic receptors, enabling rapid adaptation to stressors. This includes increased cardiac output^18^, lipolysis^19^, glycogenolysis, and gluconeogenesis^20^. Due to their short half-life, catecholamines have limited utility for assessing stress longitudinally.

Both axes share stress-induced metabolic reprogramming and an increased energy expenditure across multiple organs, including the brain, adrenal glands, heart, liver, muscle, and adipose tissue^21,22^. Metabolic total-body positron emission tomography (PET) enables simultaneous imaging of the brain and peripheral organs, offering a systems-level perspective on the metabolic response network in health and disease^23,24^.

[^18^F]fluoro-2-deoxy-D-glucose PET ([^18^F]FDG PET), traditionally used in oncology and neurology, quantifies the uptake of a radioactive glucose analog as a proxy for glucose metabolism. [^18^F]FDG PET can be combined with computed tomography or magnetic resonance imaging ([^18^F]FDG PET/CT or [^18^F]FDG PET/MRI), enabling exact anatomical allocation of glucose metabolism changes in real time in patients (dynamic PET).

PET brain imaging studies have revealed depression-associated metabolic alterations across multiple brain regions^25^. Combined with questionnaire-based assessment, it has been shown that cancer patients across different tumor types suffer from anxiety and depression^26^. In addition, increased amygdala [^18^F]FDG uptake, a biomarker of chronic stress, has been shown to predict survival of head and neck cancer patients and cardiovascular events in healthy individuals^1,5^.

Hence, we utilized metabolic PET techniques (including [^18^F]FDG and 14-(R,S)-[^18^F]fluoro-6-thia-heptadecanoic acid ([^18^F]FTHA), a fatty acid radiotracer) together with endocrine assessment and multi-omics analysis to assess the systemic effects of well-established acute, repeated acute, and surgical mouse stress models simultaneously in the whole body. Subsequently, we used image-derived network analysis to identify, describe, and visualize unique metabolic signatures in the different stress models. In this study, we identify distinct brain glucose metabolic patterns in models of acute and repeated acute stress, define a role for brown adipose tissue in stress responses independent of hypothermia, and show that glucose utilization, as a major energy source, is characterized by substrate shifts and organ-specific uptake. We further demonstrate prolonged fatty acid utilization following stress exposure, as well as potential differences in stress adaptation between cooling-induced and more severe stress paradigms, including alterations in adipose tissue volume and metabolic activity.

## Results

### Acute restraint stress leads to interscapular brown adipose tissue activation comparable to cold exposure as revealed by [^18^F]FDG µPET/CT

The adrenal gland (AG) is the primary organ responsible for catecholamine and glucocorticoid biosynthesis. As it is known that cooling stimulates norepinephrine production, we first assessed whether acute cooling induces increased glucose demand by analyzing a retrospective [^18^F]FDG PET/CT study in humans undergoing cold exposure (**Supplementary Fig. 1A**). At first, we evaluated the AG standardized uptake value (SUV) but recognized that the area surrounding the AG showed high glucose demand (**Supplementary Fig. 1B, C**). Hence, we back-translated into a preclinical PET/MRI mouse cooling model, which revealed that adjacent ventral spinal/periaortic (vsBAT), paravertebral, and perirenal brown adipose tissue depots had higher glucose demand, and the apparent AG uptake was mainly driven by spillover (**Supplementary Fig. 1D-H**). These understudied BAT depots not only appear to respond to beta-adrenergic stimulus but also might have translational implications. Thus, we continued studying mouse models across a range of stressors to investigate how BAT mediates the stress-induced metabolic response.

Hence, we next examined whether acute psychological stress, specifically restraint stress, would also affect glucose metabolism in BAT similarly to known effects of hypothermia in cold exposure in healthy male and female C57BL/6J mice (**Fig. 1A**). We found a significant twofold increase in interscapular brown adipose tissue (iBAT) [^18^F]FDG uptake after one hour of restraint comparable to the established BAT activation via acute cold exposure (3.2 vs 3.1 SUV, cooling vs restraint) (**Fig. 1B, C, Supplementary Fig. 2A**). VsBAT showed activation by increased uptake during acute cooling but not during restraint stress (1.7 vs 1.2 SUV, cooling vs restraint), revealing stress-specific BAT depot responses (**Supplementary Fig. 2B**). We also observed slight increases (+34%) in bone marrow uptake after acute cold exposure, potentially originating from bone marrow adipose tissue (BMAT) (**Supplementary Fig. 2B**).

**Figure 1:**
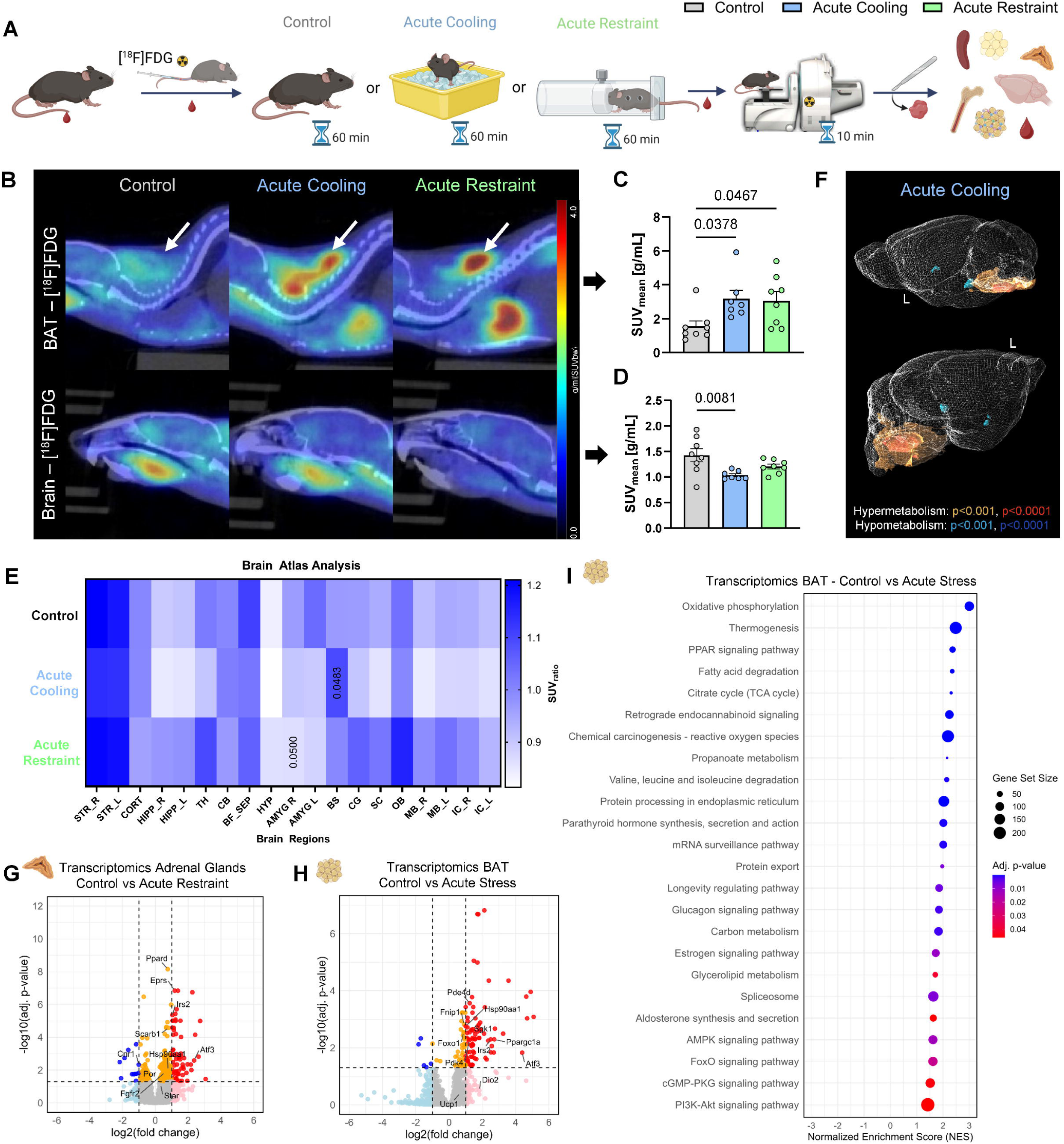
Acute restraint stress leads to interscapular brown adipose tissue activation comparable to cold exposure as revealed by [^18^F]FDG µPET/CT imaging. **A.** Illustration of the experimental setup used for the acute stress models. C57BL/6J mice were exposed to one hour of cooling (acute cooling, blue), restraint stress (acute restraint, green), or kept at control conditions (control, grey). Created in BioRender. Vraka, C. (2026) https://BioRender.com/krw61di. **B.** Representative [^18^F]FDG µPET/CT images of iBAT and brain in control, acute cooling, and acute restraint mice. **C, D.** Quantification (SUV_mean_) of [^18^F]FDG uptake via static µPET/CT imaging in iBAT (C) and brain (D) in control (n = 8), acute cooling (n = 7), and acute restraint (n = 8) mice. **E.** [^18^F]FDG uptake (SUV_ratio_) displayed as a heatmap of representative brain regions in control (n = 8), acute cooling (n = 7), and acute restraint (n = 8) mice extracted from a brain atlas (M. Mirrione) from µPET/CT images and proportionally scaled to total brain uptake. Low relative uptake is indicated in white, while high relative uptake is blue. Exact q-values of comparisons to the control group are indicated. Abbreviations of the brain regions are as follows: STR – Striatum, CORT – Cortex, HIPP – Hippocampus, TH – Thalamus, CB – Cerebellum, BF_SEP – Basal Forebrain / Septum, HYP – Hypothalamus, AMYG – Amygdala, BS – Brainstem, CG – Central Gray, SC – Superior Colliculus, OB – Olfactory Bulb, MB – Midbrain, IC – Inferior Colliculus. **F.** Voxel-wise exploratory comparison of normalized brain [^18^F]FDG uptake performed with statistical parametric mapping (SPM12) using a one-way ANOVA model with post-hoc t-contrasts and thresholds of p < 0.001 and p < 0.0001 (uncorr.) to render significantly changed areas with the PMOD 3D tool (Ver. 4.4). Images show only the differences between control (n = 8) and acute cooling (n = 7) mice. The yellow and red areas indicate hypermetabolism, while the blue and dark blue areas indicate hypometabolism. The full results are provided in Supplementary Fig. 3A. **G, H.** Bulk RNA-sequencing results of adrenal glands (G) and iBAT (H) in control (G, n = 5; H, n = 3), acute restraint (G, n = 3), and acute stress (H, n = 3; cooling and restraint) mice. Differential gene expression was analyzed using DESeq2 with two-sided Wald tests and Benjamini-Hochberg correction and illustrated as a volcano plot (downregulated: blue, downregulated (ns): light blue, upregulated: red, upregulated (ns): light red, |log2FC| ≤ 1 & p-adj. ≤ 0.05: orange, no change: grey). The headings in the Omics datasets indicate the direction of the comparison (A vs. B), with A as the control condition. **I.** KEGG pathway gene set enrichment analysis of bulk RNA-Sequencing results of iBAT (H) illustrated as a bubble plot showing normalized enrichment score (NES), adjusted p-value (adj. p-value), and gene set size. Data were analyzed using one-way ANOVA (C, D) with Dunnett’s post-hoc test and two-way ANOVA (E) with Benjamini–Hochberg FDR correction (5% across all regions and contrasts). Data are presented as individual values and as mean ± SEM, with p-values indicated as numbers above the bars.

As in the human study, the brain was not in the field of view, and we were interested in depicting the whole anatomical HPA region and activation; we next wanted to assess metabolic changes in the brain. In contrast to BAT depots, total brain uptake was reduced (-27%, acute cooling), suggesting an overall redistribution of glucose utilization from the brain towards the periphery, such as adipose tissue (**Fig. 1B, D**). Notably, blood glucose levels remained constant across our experimental groups, excluding stress-induced hyperglycemia as a potential confounder (**Supplementary Fig. 2C**). To confirm HPA-axis activation, we measured serum corticosterone levels during and after the imaging procedure. Serum corticosterone levels increased up to sixfold after one hour of stress exposure (288 ng/mL) and twofold following imaging (114 ng/mL) compared to baseline (51 ng/mL) (**Supplementary Fig. 2D**). Post-mortem corticosterone levels did not differ between the experimental groups (**Supplementary Fig. 2E**), supporting [^18^F]FDG uptake as a sensitive readout of acute stress response.

To investigate potential perfusion effects upon stress, we measured the kinetics as TACs (time-activity curves) of [^18^F]FDG in iBAT and brain using 70 min dynamic imaging (**Supplementary Fig. 2F, G**). We observed an increase (>30%) in the early phase (0-5 min) area under the curve (AUC) for both acute stress models (**Supplementary Fig. 2H**), suggesting perfusion-dependent effects potentially reflecting stress-induced increases in blood flow. Late-phase (5–60 min) AUC was increased only under restraint stress (**Supplementary Fig. 2H**). We also observed a threefold increase in metabolic rate (MR_Glu_) in the acute restraint model (10.2 MR_Glu_) compared to controls (3.1 MR_Glu_), which strongly correlated with the SUV (**Supplementary Fig. 2H, I**). Overall, the effects observed with dynamic imaging were less prominent than with static imaging, highlighting metabolic suppression via isoflurane anesthesia and unconscious tracer uptake and hence relief from stress as a crucial confounder. This was further supported by the absence of significant changes in total brain uptake or brain MR_Glu_ for both stressors, as assessed by dynamic imaging (**Supplementary Fig. 2J, K**).

As the brain is responsible for perception, processing, and signaling of the stress response, we sought to further examine the reduction in [^18^F]FDG uptake observed with static imaging by assessing regional changes. Brain atlas analysis revealed a relative increase in the brainstem (+13%) upon acute cooling and decreased relative uptake in the right amygdala (-12%) upon acute restraint (**Fig. 1E**). Voxel-based analysis confirmed hypermetabolic areas in the brainstem after acute cold exposure (**Fig. 1F, Supplementary Fig. 3A**), which aligns with thermoregulatory and sympathetic nervous system (SNS) function. Voxel-wise differences between acute restraint and control remained exploratory, whereas differences between acute cooling and acute restraint survived FDR and FWE correction (**Supplementary Fig. 3A, B**), revealing stress-specific differences that coincide with distinct adipose tissue depot activity.

To study the underlying molecular mechanisms and confirm the metabolic activation of BAT independently of glucose metabolism, we performed bulk RNA sequencing and proteomics of adrenal glands and iBAT. Adrenal glands showed significant changes in acute stress groups compared to controls, notably in genes related to steroidogenesis, such as *Scarb1, Por,* and *Star,* and in pathways such as dopaminergic synapses and cortisol/aldosterone synthesis and secretion (**Fig. 1G, Supplementary Fig. 4A-C**). Both stressors displayed a similar gene expression profile and showed no upregulation of glucose metabolism pathways (**Supplementary Fig. 4D-F**), consistent with the observed lack of [^18^F]FDG uptake increase. In iBAT, we observed marked upregulation of genes involved in mitochondrial metabolism and stress adaptation, including *Ppargc1a, Atf3, Foxo1, Fnip1, Sgk1,* and *Hsp90aa1* (**Fig. 1H**). KEGG gene set enrichment analysis revealed upregulation of oxidative phosphorylation, thermogenesis, fatty acid metabolism, amino acid utilization, and glucagon signaling pathways (**Fig. 1I, Supplementary Fig. 4G-K**), further supporting [^18^F]FDG uptake as a marker of overall BAT activity rather than solely glucose uptake.

Collectively, these findings show that acute psychological and cold stress engage a common metabolic response characterized by brown adipose tissue activation, stressor-specific regional changes in the brain, and transcriptional changes, highlighting metabolic imaging as a sensitive and translational readout of acute stress response.

### Repeated acute stress exposure reveals differences in the metabolic responses to cooling and restraint stress

Next, we aimed to study the effects of repeated acute stress exposure to investigate potential adaptation and remodeling mechanisms in our target organs. Mice were subjected to the same stressors as in the acute models, but for 1 hour daily over 1 week, with a 24-hour recovery period before imaging (**Fig. 2A**).

**Figure 2:**
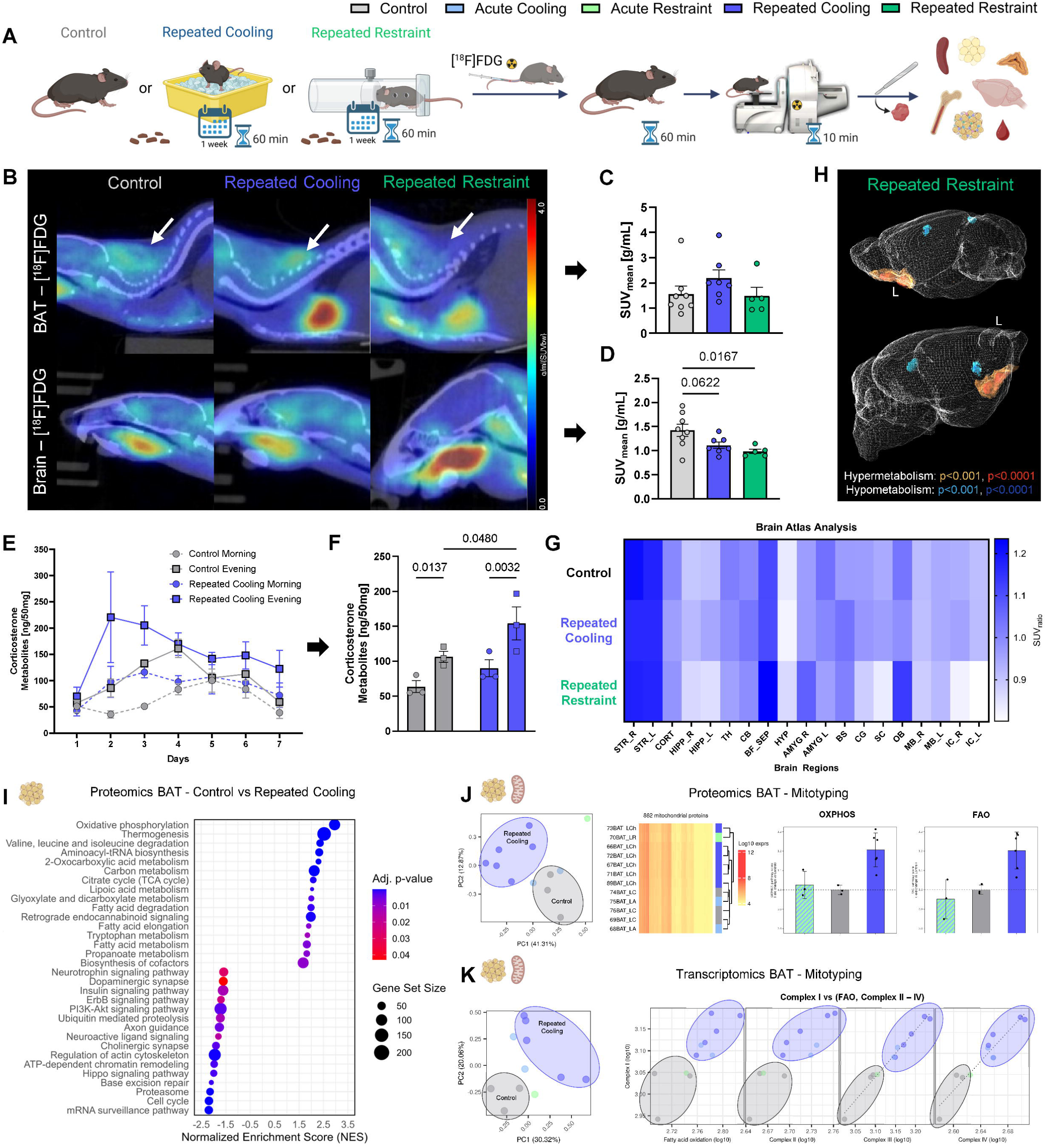
Repeated acute stress reveals differences in the metabolic responses to cooling and restraint stress. **A.** Illustration of the experimental setup of the repeated acute stress models. C57BL/6J mice were exposed to one hour of cooling (repeated cooling, blue) or restraint stress (repeated restraint, green) daily over one week, or kept at control conditions (control, grey). Created in BioRender. Vraka, C. (2025) https://BioRender.com/4qwwwa5. **B.** Representative [^18^F]FDG µPET/CT images of iBAT and brain in control, repeated cooling, and repeated restraint mice. **C, D**. Quantification (SUV_mean_) of [^18^F]FDG uptake via static µPET/CT imaging in iBAT (**C**) and brain (**D**) in control (n = 8), repeated cooling (n = 7), and repeated restraint (n = 5) mice. **E, F**. Levels of fecal corticosterone metabolites (ng/50 mg) sampled throughout one week in the morning and evening in control (n = 3) and repeated cooling (n = 3) mice. Illustrated as daily values over the course of one week (**E**) and as mean morning and evening levels per mouse across the week (**F**). **G.** [^18^F]FDG uptake (SUV_ratio_) displayed as a heatmap of representative brain regions in control (n = 8), repeated cooling (n = 7), and repeated restraint (n = 5) mice extracted from a brain atlas (M. Mirrione) from µPET/CT images and proportionally scaled to total brain uptake. Low relative uptake is indicated in white, while high relative uptake is blue. Exact q-values of comparisons to the control group are indicated. Abbreviations of the brain regions are as follows: STR – Striatum, CORT – Cortex, HIPP – Hippocampus, TH – Thalamus, CB – Cerebellum, BF_SEP – Basal Forebrain / Septum, HYP – Hypothalamus, AMYG – Amygdala, BS – Brainstem, CG – Central Gray, SC – Superior Colliculus, OB – Olfactory Bulb, MB – Midbrain, IC – Inferior Colliculus. **H.** Voxel-wise exploratory comparison of normalized brain [^18^F]FDG uptake performed with statistical parametric mapping (SPM12) using a one-way ANOVA model with post-hoc t-contrasts and thresholds of p < 0.001 and p < 0.0001 (uncorr.) to render significantly changed areas with the PMOD 3D tool (Ver. 4.4). Images show only the differences between control (n = 8) and repeated restraint (n = 5) mice. The yellow and red areas indicate hypermetabolism, while the blue and dark blue areas indicate hypometabolism. The full results are provided in **Supplementary Fig. 6A**. **I.** KEGG pathway gene set enrichment analysis of Proteomics results analyzed using LIMMA with two-sided moderated t-tests and Benjamini-Hochberg correction of iBAT in control (n = 3) and repeated cooling (n = 6) mice illustrated as a bubble plot showing normalized enrichment score (NES), adjusted p-value (adj. p-value), and gene set size. The headings in the Omics datasets indicate the direction of the comparison (A vs. B), with A as the control condition. **J.** Mitotyping analysis of Proteomics results in iBAT of control (n = 3), acute stress (n = 3; cooling and restraint), and repeated cooling (n = 6) mice. Left to right: 1. PCA analysis on mitochondrial Pathway Priority Scores (mitoPPS) displaying PC1–PC2 projections visualizing the major axes of variation in mitochondrial pathway prioritization. 2. Hierarchical clustering of samples according to 862 mitochondrial proteins. 3, 4. Oxidative phosphorylation (OXPHOS, 3.) and fatty acid oxidation (FAO, 4.) mitochondrial pathway scores per experimental group. **K.** Mitotyping analysis of Transcriptomics results in iBAT of control (n = 3), acute stress (n = 3; cooling and restraint), and repeated cooling (n = 5) mice. Left to right: 1. PCA analysis on mitochondrial Pathway Priority Scores (mitoPPS) displaying PC1–PC2 projections visualizing the major axes of variation in mitochondrial pathway prioritization. 2. Plots showing OXPHOS (Complex I-IV) and FAO expression profiles of samples visualizing the molecular specialization of mitochondria. Data were analyzed using one-way ANOVA (**C, D**) with Dunnett’s post-hoc test, two-way ANOVA (**G**) with Benjamini–Hochberg FDR correction (5% across all regions and contrasts), and two-way repeated-measures ANOVA (**F**) with Fisher’s LSD post-hoc test. Data are presented as individual values and as mean ± SEM, with p-values indicated as numbers above the bars. Pictograms of organs in I-K were taken from *BioRender. Vraka, C. (2026)* https://BioRender.com/d1uqx40.

In contrast to the acute model, repeated restraint stress did not show prolonged iBAT activation, while repeated cooling still showed a trend towards increased iBAT [^18^F]FDG uptake (2.2 SUV) and significance for vsBAT (1.6 SUV) (**Fig. 2B, C, Supplementary Fig. 5A),** illustrating divergent BAT depot activation upon repeated acute stress exposure. Skeletal muscle tissue uptake was also found to be increased twofold by repeated cooling but not repeated restraint (**Supplementary Fig. 5A**), indicating muscle activity from continuous shivering and increased movement during cooling exposure. Total brain uptake was significantly decreased (-30%) in the repeated restraint model, with a trend (-22%) for repeated cooling (**Fig. 2B, D**). Unlike in the acute restraint model, this decrease was not accompanied by compensatory increases in uptake in other analyzed organs (**Supplementary Fig. 5A**).

Also, for the repeated acute stress models, blood glucose levels were constant across all groups (**Supplementary Fig. 5B**). On the other hand, serum corticosterone levels were elevated in repeated cooling (168 ng/mL) but not for repeated restraint (124 ng/mL), indicating stressor-specific endocrine responses possibly due to dysregulation or habituation (**Supplementary Fig. 5C**). Non-invasive fecal corticosterone metabolite analysis further revealed expected circadian-dependent variations in HPA axis activity and showed peaking corticosterone release after two to three days of repeated cooling (205 ng/50mg) compared to controls (133 ng/50mg) (**Fig. 2E, F)**. Repeated cooling additionally led to a reduction (-14%) in body weight compared to controls and repeated restraint without altering overall body composition (**Supplementary Fig. 5D, E**), which may be explained by the observed increased metabolic activation, but could also be caused by decreased food intake, which was not assessed in this study. Taken together, these findings demonstrate that repeated cooling, unlike repeated restraint, increases corticosterone secretion and BAT tissue adaptation, resulting in prolonged metabolic activation and potential weight loss.

Overall decrease in total brain uptake was also consistent in the repeated acute stress groups (-31%, repeated restraint), while regional changes in brain uptake were less pronounced than in the acute stress models. Brain atlas analysis showed no significant differences in regional brain uptake for both repeated acute stress models (**Fig. 2G**). Voxel-wise analysis only revealed a hypermetabolic cluster for the repeated restraint group in the forebrain and olfactory bulb area (**Fig. 2H)** and a hypometabolic cluster between repeated cooling and repeated restraint. The other findings remained exploratory, highlighting only minor stressor-specific differences (**Supplementary Fig. 6A, B**). Together, this suggests that regional brain metabolic changes are less pronounced after repeated stress and a 24 h recovery period, while alterations in systemic glucose metabolism persist.

We sought to further investigate the immunometabolic response to corticosterone, a known immunosuppressant, since we identified changes in [^18^F]FDG uptake in bone marrow after acute cooling (**Supplementary Fig. 2B**). For this, we performed RadioFlow analysis in spleen and bone marrow, which measures [^18^F]FDG uptake in sorted cell populations after *in vivo* tracer distribution (**Supplementary Fig. 7A**). We found that indeed repeated cooling showed trends toward a reduction (-15%) in splenic B-cell numbers and a relative reduction (-70%) in [¹□F]FDG uptake in bone marrow T cells compared to controls (**Supplementary Fig. 7B–E**), suggesting altered immune activity.

Consistent with this, we observed downregulation of pro-inflammatory immune signaling pathways, including TNF and B cell receptor signaling in iBAT in our repeated cooling model **(Supplementary Fig. 8A)**. Further results in iBAT primarily revealed increased expression of mitochondrial genes in the repeated cooling group compared to the controls (**Supplementary Fig. 8B**). KEGG pathway analysis largely matched the acute stress models (**Supplementary Fig. 8A, C-H**), but with reduced tyrosine, cysteine, methionine, and arachidonic acid metabolism in our acute stress models compared to repeated cooling indicating a slightly shifted metabolism towards amino acids. Proteomic profiling confirmed enhanced mitochondrial and metabolic activity in iBAT, alongside downregulation of signaling and cell cycle pathways (**Fig. 2I, Supplementary Fig. 8I**). Together, these results confirm prolonged metabolic activation and adaptation of iBAT following repeated cold exposure of just one hour per day.

RNA-seq and proteomic analysis of adrenal glands predominantly indicated downregulation of metabolic, hormone, and signaling pathways, including glycerophospholipid metabolism, thyroid hormone signaling, and dopaminergic synapse pathways, after repeated cooling (**Supplementary Fig. 8J-M)**. Overall, proteomics revealed more extensive changes in our repeated acute models compared to the acute stress models, whereas transcriptomic changes were more pronounced in the acute models, indicating time-dependent adaptation at both the protein and transcriptional levels. Based on our Omics data from acute models, we performed RT-qPCR on selected genes in iBAT and found significant upregulation of *Crebbp* and *Sgk1* in the repeated restraint model, with trends toward upregulation of *Adrb3* and *Fabp4* (**Supplementary Fig. 9A**), highlighting enhanced beta-adrenergic signaling and lipid metabolic adaptation in iBAT.

To further investigate mitochondrial adaption, we employed a novel Mitotyping analysis^27,28^ of transcriptomics and proteomics data from our stress models. We identified pronounced changes in mitochondrial gene and protein expression in iBAT, particularly following repeated cooling. These changes are characterized by increased OXPHOS and fatty acid oxidation (FAO) pathways and distinct clustering of samples based on principal component analysis (PCA) of mitochondrial Pathway Priority Scores (mitoPPS) and hierarchical clustering analysis (**Fig. 2 J, K**). These findings indicate mitochondrial metabolic remodeling in iBAT in response to repeated cooling, consistent with the increased energetic demands associated with this stress paradigm. Inspection of the corresponding PCA loadings identified OXPHOS and metabolic pathways contributing most strongly to the major axes of variation (**Supplementary Fig. 9B**). In contrast to iBAT, adrenal glands showed less distinct clustering in PCA and hierarchical analysis, together with fewer changes in mitochondrial pathway signatures, further supporting iBAT as a metabolically responsive tissue during repeated stress exposure, including at the mitochondrial level (**Supplementary Fig. 9C-E**).

Overall, repeated acute stress exposure reveals a clear difference between stress modalities, with repeated cooling driving sustained BAT metabolic activity, endocrine response, and mitochondrial remodeling, whereas repeated restraint shows marked habituation or blunting. These results indicate that metabolic imaging sensitively reflects stressor-specific adaptation and remodeling beyond the acute response.

### Surgery and anesthesia induce severe, prolonged metabolic changes in brain and adipose tissue metabolism

To implement a more severe and translational stress model, we next sought to investigate surgical procedures using longitudinal [^18^F]FDG PET to track metabolic changes over 10 days (**Fig. 3A**). To account for the effects of the anesthetic-analgesic regimen on hypothalamic thermoregulation, we included an additional anesthesia group and monitored body temperature.

**Figure 3:**
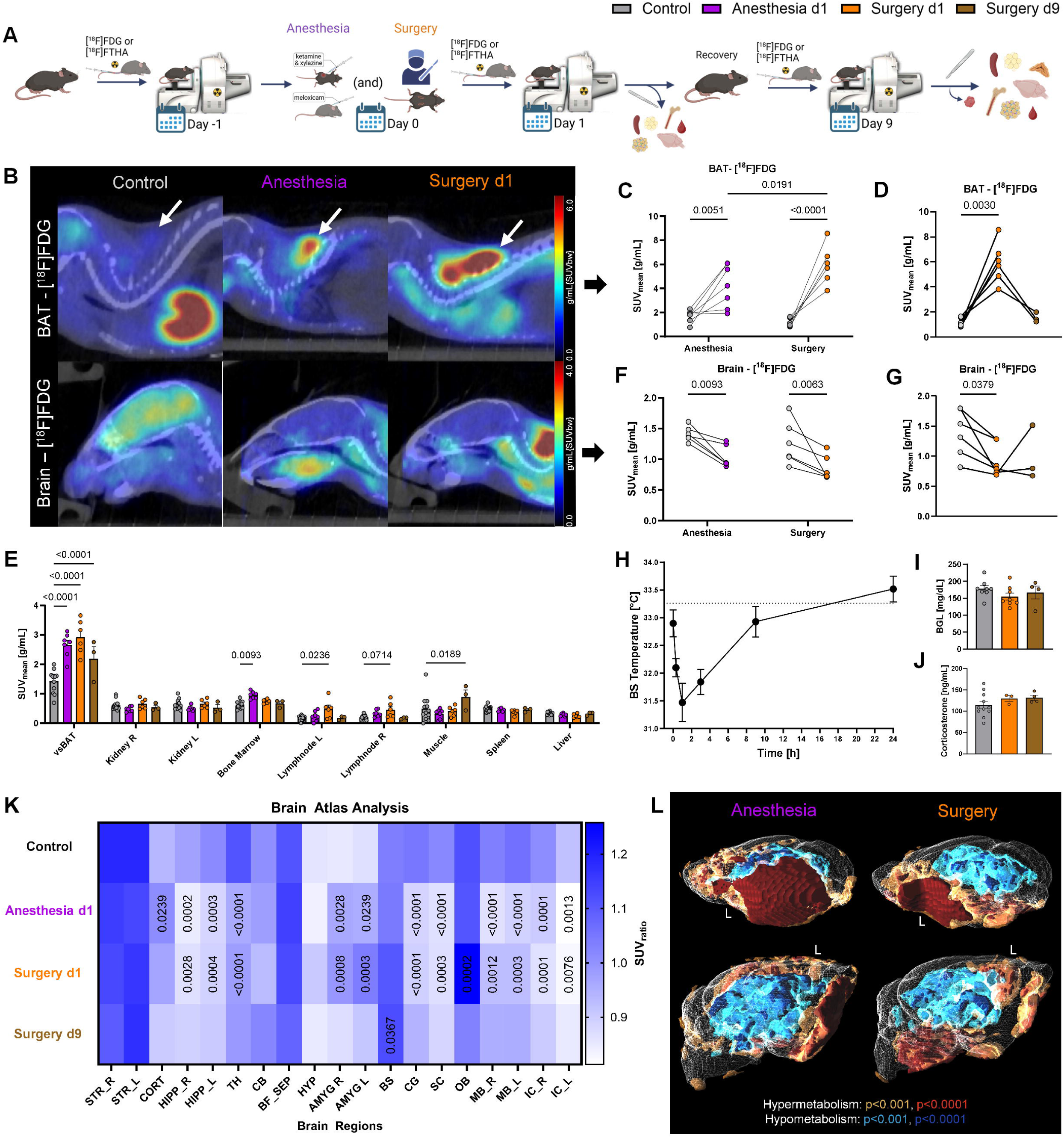
Surgery and anesthesia induce severe, prolonged metabolic changes in brain and adipose tissue metabolism. **A**. Illustration of the experimental setup used for the surgery models. C57BL/6J mice underwent baseline scans (control, grey) followed by either the anesthetic/analgesic regimen (anesthesia d1, purple) or a surgical procedure (surgery d1, orange), including a scan after 1 day, and a follow-up scan after 9 days (surgery d9, brown). Created in BioRender. Vraka, C. (2026) https://BioRender.com/cgbr7f2. **B**. Representative [^18^F]FDG µPET/CT images of iBAT and the brain in control, anesthesia d1, and surgery d1 mice. **C, D**. Quantification (SUV_mean_) of [^18^F]FDG uptake via static µPET/CT imaging in iBAT in control (n = 7) and anesthesia d1 (n = 7) mice, control (n = 6) and surgery d1 (n = 6) mice (**C**), and in control (n = 6), surgery d1 (n = 6), and surgery d9 (n = 3) mice (**D**). **E**. Quantification (SUV_mean_) of [^18^F]FDG uptake via static µPET/CT imaging in other tissues in control (n = 12), anesthesia d1 (n = 7), surgery d1 (n = 6), and surgery d9 (n = 3) mice. **F, G**. Quantification (SUV_mean_) of [^18^F]FDG uptake via static µPET/CT imaging in brain in control (n = 6) and anesthesia d1 (n = 6) mice, control (n = 6) and surgery d1 (n = 6) mice (**F**), and in control (n = 6), surgery d1 (n = 6) and surgery d9 (n = 3) mice (**G**). **H.** Body surface temperature in anesthesia (n = 6-7) mice post xylazine/ketamine anesthesia measured with an infrared thermometer. **I.** Blood glucose levels in control (n = 8), surgery d1 (n = 8), and surgery d9 (n = 4) mice. **J.** Serum corticosterone levels in control (n = 11), surgery d1 (n = 3), and surgery d9 (n = 4) mice. **K.** [^18^F]FDG uptake (SUV_ratio_) displayed as a heatmap of representative brain regions in control (n = 12), anesthesia d1 (n = 6), surgery d1 (n = 6), and surgery d9 (n = 3) mice extracted from a brain atlas (M. Mirrione) from µPET/CT images and proportionally scaled to total brain uptake. Low relative uptake is indicated in white, while high relative uptake is blue. Exact q-values of comparisons to the control group are indicated. Abbreviations of the brain regions are as follows: STR – Striatum, CORT – Cortex, HIPP – Hippocampus, TH – Thalamus, CB – Cerebellum, BF_SEP – Basal Forebrain / Septum, HYP – Hypothalamus, AMYG – Amygdala, BS – Brainstem, CG – Central Gray, SC – Superior Colliculus, OB – Olfactory Bulb, MB – Midbrain, IC – Inferior Colliculus. **L.** Voxel-wise exploratory comparison of normalized brain [^18^F]FDG uptake performed with statistical parametric mapping (SPM12) using a one-way ANOVA model with post-hoc t-contrasts and thresholds of p < 0.001 and p < 0.0001 (uncorr.) to render significantly changed areas with the PMOD 3D tool (Ver. 4.4). Images show only the differences between control (n = 12) and anesthesia d1 (n = 6, left) as well as control (n = 12) and surgery d1 (n = 6, right). The yellow and red areas indicate hypermetabolism, while the blue and dark blue areas indicate hypometabolism. The full results are provided in **Supplementary Fig. 10A**. Data were analyzed using two-way repeated-measures ANOVA (**C, F**) with Fisher’s LSD post-hoc test, mixed-effects analysis (**D, G**) with Dunnett’s post-hoc test, two-way ANOVA (**E, K**) with Dunnett’s post-hoc test (**E**) or Benjamini–Hochberg FDR correction (5% across all regions and contrasts; **K**), and one-way ANOVA (**I, J**) with Tukey’s post-hoc test. Data are presented as individual values and as mean ± SEM, with p-values indicated as numbers above the bars.

iBAT [^18^F]FDG uptake increased fivefold 24 hours after surgery and anesthesia, thereby exceeding the changes observed in acute and repeated acute stress models (**Fig. 3B, C**). The magnitude of the metabolic response differed significantly between surgery and anesthesia (6.0 SUV vs 4.2 SUV), indicating cumulative effects induced by surgical trauma on BAT activity (**Fig. 3B, C**). On day 9, iBAT uptake returned nearly to control levels (1.6 SUV vs 1.2 SUV), whereas vsBAT uptake remained elevated (2.2 SUV) (**Fig. 3D, E**). Overall, uptake increased up to twofold in vsBAT, whereas it was fivefold in iBAT, suggesting heterogeneity in the metabolic response of BAT depots. We also observed differences in uptake in the bone marrow (+50%, anesthesia d1), lymph nodes (+167%, surgery d1), and muscle (+81%, surgery d9), indicating potential influence on the immune system and increased mobility after 9 days (**Fig. 3E**). Also in the surgery model, total brain [^18^F]FDG uptake decreased (-32%, surgery d1; -27%, anesthesia d1) at 24 hours and returned to control levels at day 9 (**Fig. 3B, F, G**). We found that the animal’s body surface temperature dropped post-anesthesia (31.5°C) and returned to control levels (32.9°C) within 8 hours, suggesting that the observed increase in BAT activity 24 hours after anesthesia represents a prolonged metabolic adaptation to anesthesia that cannot be explained by acute adaptation to body temperature changes alone (**Fig. 3H**). Additionally, consistent with acute stress models, we observed no difference in blood glucose (**Fig. 3I**) or corticosterone levels (**Fig. 3J**) at 24 hours or 9 days post-surgery.

Additional brain atlas analysis revealed significant changes in regional brain uptake. We found the cortex (+8%) and amygdala (+10%) to be hypermetabolic, and the hippocampus (-12%), thalamus (-15%), central gray (-16%), superior colliculi (-14%), midbrain (-16%), and inferior colliculi (-12%) to show hypometabolism under our anesthetic-analgesic regimen. After surgery, we observed elevated relative uptake in the amygdala (+13%) and olfactory bulb (+11%) and decreased uptake in the hippocampus (-11%), thalamus (-12%), central gray (-13%), superior colliculi (-12%), midbrain (-11%), and inferior colliculi (-11%), matching most of the regional differences seen after anesthesia (**Fig. 3K**). Voxel-wise analysis confirmed these changes and further illustrated large hypo- and hypermetabolic areas across these regions (**Fig. 3L, Supplementary Fig. 10A**). Differences between surgery and anesthesia are as expected, less pronounced, but surgery shows hypermetabolic areas around the brainstem and hypothalamus and hypometabolic areas around the cortex (**Supplementary Fig. 10A, B**). Consistent with total brain uptake, regional uptake reaches control levels after 9 days except for an elevated relative uptake in the brainstem (**Fig. 3K**), reflecting recovery of brain metabolism.

Taken together, these results show that surgery and anesthesia elicit a severe, prolonged metabolic stress response that exceeds that of our non-invasive stress models, characterized by high BAT activation, widespread effects on peripheral organs, and altered brain glucose utilization, which are largely reversible within 9 days.

### Metabolic reprogramming of adipose tissue depots after surgery and anesthesia reveals significant systemic effects of surgical interventions

Unlike in the acute and repeated acute stress models, we found a twofold increase in [^18^F]FDG uptake and a decrease (-20%) in volume in inguinal white adipose tissue (iWAT) after anesthesia and surgery (**Fig. 4A-C**). White adipose tissue, such as iWAT, can adapt to metabolic challenges by whitening or browning and increasing or decreasing its thermogenic function. Hence, due to these remodeling properties of adipose tissue, we wanted to further investigate fatty acid metabolism by using 14-(R, S)-[^18^F]fluoro-6-thia-heptadecanoic acid ([^18^F]FTHA), a free fatty acid analog. In iWAT, [^18^F]FTHA uptake decreased on days 1 and 9 (-18% and -34%), which contrasts with [^18^F]FDG, suggesting a shift from lipid utilization to glucose consumption after surgery, potentially reflecting enhanced lipolysis and tissue remodeling (**Fig. 4D, E**). In iBAT, [^18^F]FTHA showed increased (+45%) free fatty acid uptake 24 hours post-surgery, indicating enhanced β-oxidation. Notably, [^18^F]FTHA uptake remained elevated on day 9 in two animals, suggesting sustained lipid utilization, but the group-level difference remained non-significant. However, the magnitude relative to controls was nine times lower than that of [^18^F]FDG, suggesting that glucose is the predominant energy source, ensuring rapid energy supply (**Fig. 4D, F**).

**Figure 4:**
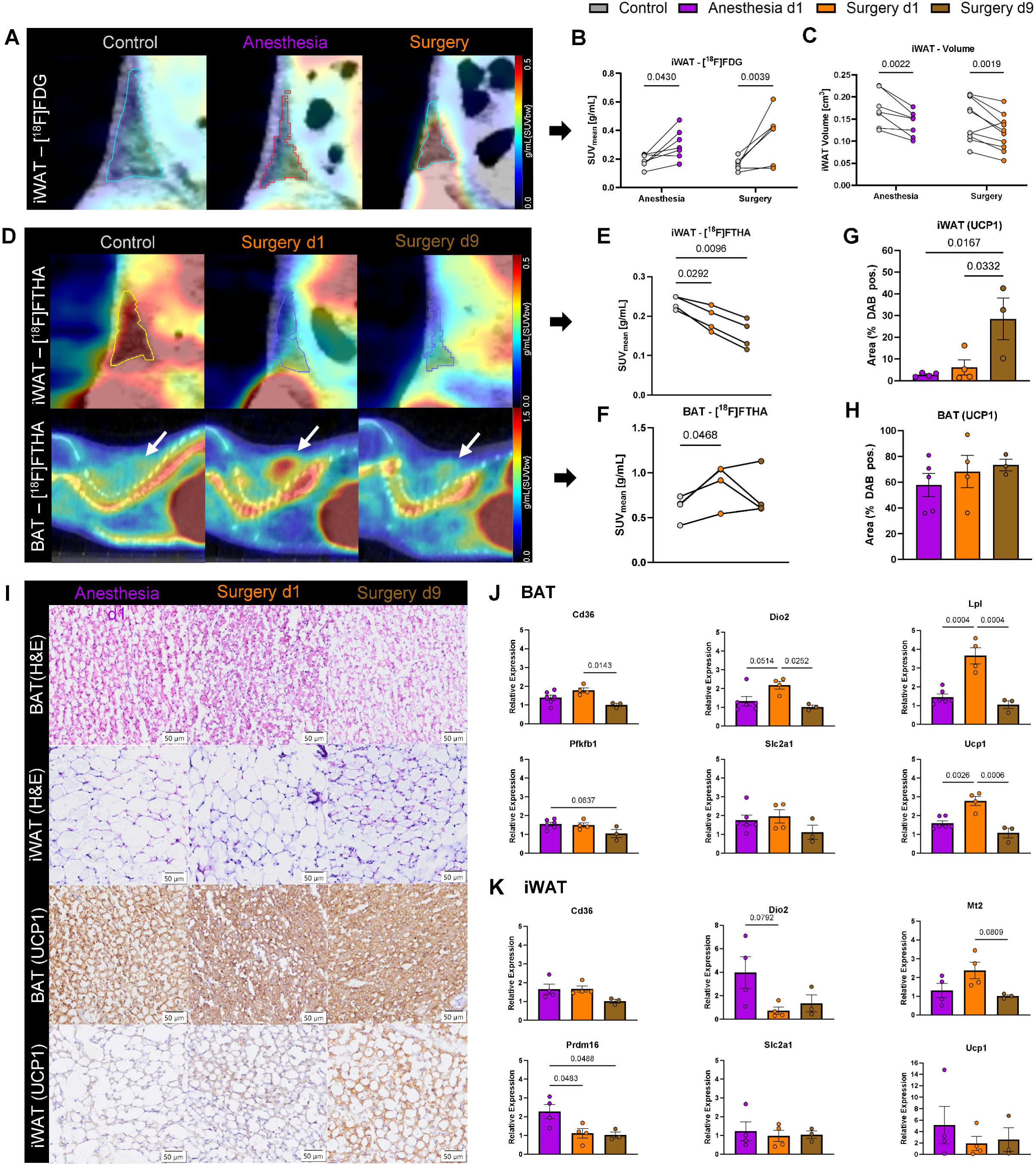
Metabolic reprogramming of adipose tissue depots after surgery and anesthesia reveals significant systemic effects of surgical interventions. **A.** Representative [^18^F]FDG µPET/CT images of iWAT in control, anesthesia d1, and surgery d1 mice. **B.** Quantification (SUV_mean_) of [^18^F]FDG uptake via static µPET/CT imaging in iWAT in control (n = 7) and anesthesia d1 (n = 7) mice, and control (n = 6) and surgery d1 (n = 6) mice. **C.** Quantification of iWAT volume (cm^3^) in control (n = 7) and anesthesia d1 (n = 7) mice, and control (n = 10) and surgery d1 (n = 10) mice. **D.** Representative [^18^F]FTHA µPET/CT images of iWAT and iBAT in control, surgery d1, and surgery d9 mice. **E, F**. Quantification (SUV_mean_) of [^18^F]FTHA uptake via static µPET/CT imaging in iWAT (**E**) and iBAT (**F**) in control (n = 4), surgery d1 (n = 4), and surgery d9 (n = 4) mice. **G, H**. Quantification of DAB-positive areas (UCP1) of iWAT (**G**) and iBAT (**H**) in anesthesia d1 (**G**, n = 4; **H**, n = 5), surgery d1 (n = 4), and surgery d9 (n = 3) mice. **I**. Representative H&E and DAB (UCP1) staining of iWAT and iBAT in anesthesia d1, surgery d1, and surgery d9 mice (scale bars 50 μm). **J, K**. Relative gene expression of iBAT (**J**) and iWAT (**K**) tissue assessed via RT-qPCR in anesthesia d1 (**J**, n = 6; **K**, n = 4), surgery d1 (n = 4), and surgery d9 (n = 3) mice quantified via the ΔΔCt method. Data were analyzed using two-way repeated-measures ANOVA (**B, C**) with Fisher’s LSD post-hoc test, one-way repeated-measures ANOVA (**E, F**) with Dunnett’s post-hoc test, and one-way ANOVA (**G, H, J, K**) with Tukey’s post-hoc test. Data are presented as individual values and as mean ± SEM, with p-values indicated as numbers above the bars.

To observe adipose tissue remodeling at the molecular level, we assessed UCP1 protein expression by IHC, which showed four- to tenfold elevated UCP1 expression in iWAT on day 9, confirming adipose tissue browning post-surgery. iBAT showed only a slight trend towards increased UCP1 expression (**Fig. 4G-I**). Additional RT-qPCR analysis revealed transcriptional upregulation of markers in iBAT and iWAT associated with browning (*Prdm16*), thermogenesis (*Dio2, Ucp1*), lipid metabolism (*Cd36, Lpl*), and sympathetic innervation (*Mt2*) post-surgery. The differences in protein and transcriptional expression in iWAT likely reflect post-transcriptional regulation and delayed accumulation of UCP1 during adipose tissue remodeling. The concomitant reduction in iWAT volume, together with reduced lipid uptake and increased glucose uptake, is consistent with enhanced lipolysis and a metabolic shift away from lipid storage toward substrate oxidation, rather than transcriptional control of Ucp1 expression. Interestingly, the expression of *Dio2* (+64%), *Lpl* (+151%), and *Ucp1* (+74%) in iBAT was markedly increased after surgery compared with the anesthesia group, highlighting the effects of surgical trauma beyond thermogenic activation, which we can depict minimally invasively, with increased [^18^F]FDG uptake (**Fig. 4J, K**, **Supplementary Fig. 11A, B**).

In summary, surgery and sedative–analgesic anesthetic regimens drive remodeling across adipose tissue depots, characterized by browning and metabolic reprogramming of iWAT and iBAT, and a shift in fuel utilization that extends beyond thermogenesis, underscoring the systemic impact and severity of surgical stress.

### Network analysis gives insight into the systemic metabolic fingerprint of different stress models

To investigate central and peripheral organ crosstalk and regional brain changes, we sought to determine whether network analysis can elucidate stress-specific metabolic patterns. To investigate the complete metabolic interplay across adipose tissue and brain regions, a partial-correlation network model was used, in which network structures were compared with their corresponding weight matrices. High network correlations reflect similar metabolic coordination across various adipose tissue depots and brain regions. In all stress models, the interaction between brain and vsBAT was lost, indicating a loss of correlation as an overall stress indicator (**Fig. 5A**). Across the stress models, the brain correlated with iWAT, iBAT (**Fig. 5 B, C**), or both, highlighting increased communication with these adipose tissue depots. The correlation between iWAT and iBAT shifted from negative to positive in our acute stress model but was lost entirely in repeated acute stress and surgical intervention, possibly due to remodeling of these adipose depots under prolonged stress exposure.

**Figure 5:**
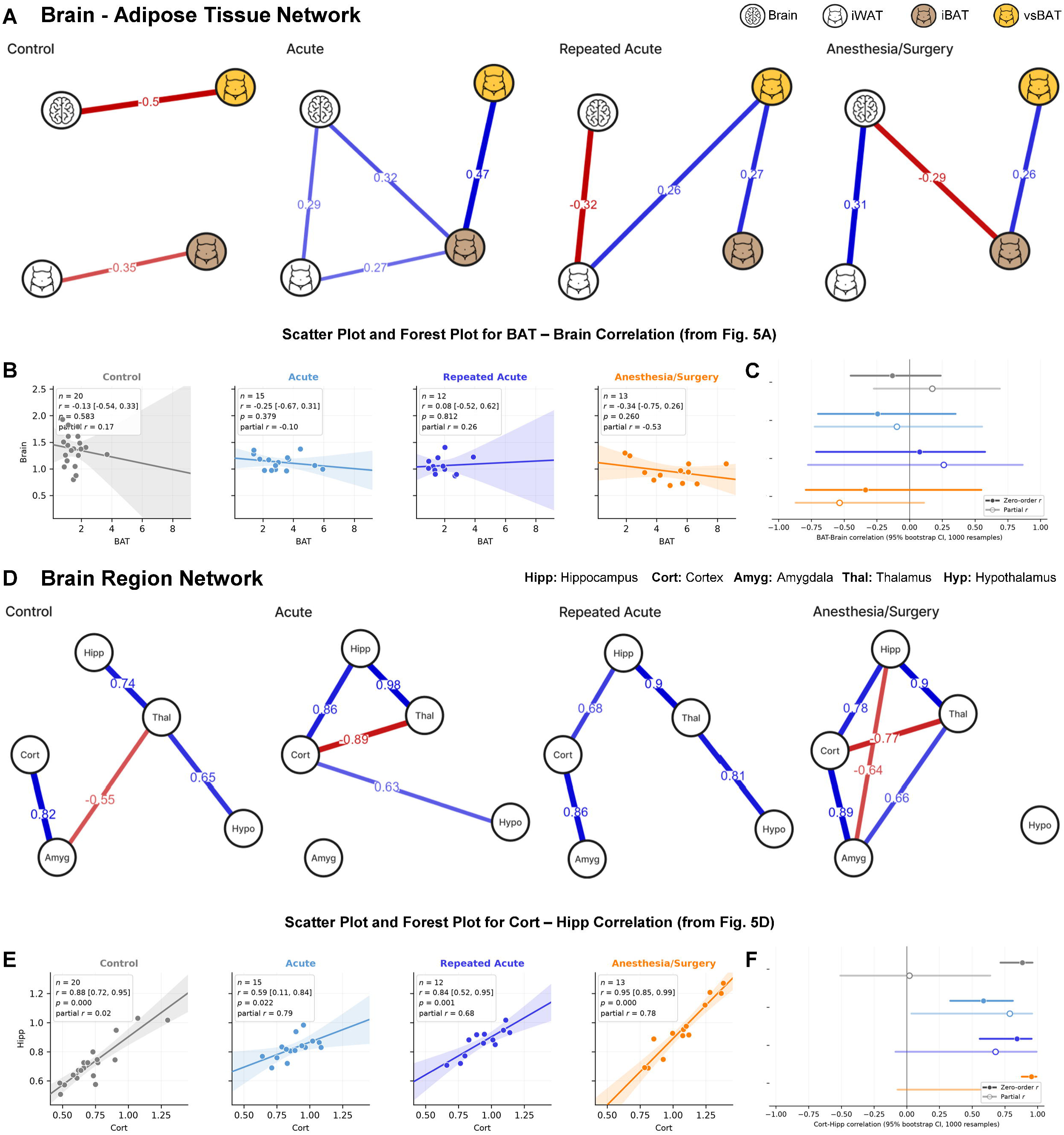
Network analysis gives insight into the systemic metabolic fingerprint of different stress models. **A.** Partial correlation network analysis of the brain and adipose tissue depots based on [^18^F]FDG PET imaging data (SUV_mean_). Networks are shown for control (n = 20), acute (n = 15), repeated acute (n = 12), and anesthesia/surgery (n = 13) groups. Only significant partial correlations with a correlation coefficient of |r| > 0.25 are displayed. **B, C**. Scatter plots (**B**) illustrating the correlation between total brain and iBAT [^18^F]FDG uptake (SUV_mean_) for each experimental group from **A** and a forest plot (**C**) showing each group’s zero-order correlation coefficient (filled symbols) and partial correlation coefficient (empty symbols), with their bootstrap estimates (95% confidence intervals, 1000 resamples). Shaded bands indicate 95% confidence intervals of the fitted regression lines. **D**. Partial correlation network analysis of stress-related brain regions based on [^18^F]FDG PET imaging data (SUV_ratio_) scaled to total brain uptake. Networks are shown for control (n = 20), acute (n = 15), repeated acute (n = 12), and anesthesia/surgery (n = 13) groups. Only significant partial correlations with a correlation coefficient of |r| > 0.25 are displayed. **E, F**. Scatter plots (**E**) illustrating the correlation between cortex (cort) and hippocampus (hipp) uptake (SUV_ratio_) for each experimental group from **D** and a forest plot (**F**) showing each group’s zero-order correlation coefficient (filled symbols) and partial correlation coefficient (empty symbols), with their bootstrap estimates (95% confidence intervals, 1000 resamples). Shaded bands indicate 95% confidence intervals of the fitted regression lines. Correlations (edges) are displayed as positive (blue), negative (red), magnitude (thickness), strength (color intensity), and their coefficient value.

In brain region networks (**Fig. 5D**), we observed a stable correlation between the hippocampus and thalamus (r = 0.74–0.98, p < 0.05) across all experimental groups. However, in the control network, the hippocampus did not correlate with the cortex, indicating a unique and stress-specific interaction observed across all stress-model networks (r = 0.68–0.86, p < 0.05) (**Fig. 5E, F**). A loss of cortex–amygdala correlation (r = 0.82–0.89, p < 0.05) was observed only in acute stress models, indicating differences between acute and repeated acute stress. Notably, the amygdala for acute stress and the hypothalamus for surgery/anesthesia appeared metabolically isolated (i.e., no correlations), suggesting distinct roles in their respective stress models. Visually, all brain networks exhibit a stressor-specific structure, highlighting this method as a distinct approach for revealing regional interactions in brain metabolism. Data points for all correlations used in network analysis are displayed in the supplementary figures (**Supplementary Fig. 12A-J**, **Supplementary Fig. 13A-N**).

In summary, our network analysis reveals multiple inter-organ correlations between the brain and adipose tissue depots, as well as across brain regions, in response to acute and repeated acute stress, visualizing stress-induced metabolic communication and remodeling.

## Discussion

Stress elicits metabolic adaptations that raise energy expenditure and trigger tissue remodeling that depend on stressor type and duration, yet objective markers remain limited, especially on a whole-body level. Advances in total-body PET, together with AI-based computational methods, have extended the use of PET imaging beyond traditional applications: it enables minimally-invasive assessment of central and peripheral organ metabolism, shifting utilization of [^18^F]FDG PET for a more complex understanding of how the entire body is affected by stress, adding new options as the exclusive established imaging marker with prognostic relevance^23–26^. We used established stress models and showed that acute, repeated acute, and surgical stress induce shared and stressor-specific metabolic signatures across brain regions and organs, particularly in adipose tissue.

Mechanistically, stress-induced BAT activation is mediated by common beta-adrenergic signaling via the sympathetic-adrenal medullary (SAM) axis^29–31^. In accordance with our data, psychological stress drives non-shivering thermogenesis (NST) in BAT via a hypothalamomedullary glutamatergic pathway^32^, and sympathetic projections can regulate thermogenesis and glucose tolerance independently in BAT^33^. To the best of our knowledge, we are the first utilizing [^18^F]FDG PET to show that psychological (restraint) and surgical stress robustly activate adipose tissue metabolism, comparable to cold exposure but independent of hypothermia, identifying brown adipose tissue (BAT) as a central stress-responsive metabolic hub. Notably, the physiological relevance of BAT in human adults was also discovered via [^18^F]FDG PET, revealing metabolically active depots in the supraclavicular and paravertebral regions during cold stimulation^34,35^. Building upon this, we show that stress-induced metabolic activity is predominantly localized to BAT, including non-classical depots such as ventral spinal BAT (vsBAT), highlighting functional heterogeneity among adipose tissue depots with translational relevance^36^. Hitherto, studies have largely focused on interscapular BAT (iBAT) and its primary function, NST, in the context of cold exposure^37,38^. Importantly, BAT activation is not uniform across stress paradigms. We show that repeated restraint stress leads to a loss of metabolic response when compared to cold exposure, as this stress form is prone to habituation, blunting the response to subsequent stress^39^. This aligns with other studies indicating that repeated restraint stress diminishes the hyperthermia response to cold exposure^40^, more so than repeated cooling alone^30^. This is partially explained by the hypothalamic-pituitary-adrenal (HPA) axis, which modulates BAT activity rapidly via GC. They primarily act as inhibitors, but depending on the species, dosage, and duration, they can also act as agonists^41,42^. However, as we did not find a correlation between GC levels and BAT activity, this suggests that endocrine output is insufficient to explain peripheral organ metabolism, and further studies are required to elucidate the exact mechanisms by which the SAM and HPA axis modulate BAT and peripheral tissue activity. Indeed, glucose uptake measured by [^18^F]FDG PET emerged as the most sensitive indicator of BAT activation across our models, providing a minimally invasive approach to assess stress and study different BAT depots simultaneously, beyond cold-induced thermogenesis. In comparison, serum GC levels were less reliable indicators of stress exposure, as sampling time (circadian rhythm) and frequency are major confounders, and continuous blood sampling itself can increase inflammation^16^. Instead, fecal GC metabolite assessment should be favored for repeated acute stress models^43^, but still requires repeated sampling and captures systemic endocrine effects, rather than organ-specific effects. Taken together, our findings support a model in which stress-induced metabolic activation is primarily driven by neural and local tissue-level processes and stable biomarker for stress rather than systemic hormone levels alone.

Across our stress models, surgical stress induced the strongest metabolic activation, accompanied by adipose tissue remodeling and upregulation of genes involved in lipid uptake and thermogenesis, supporting a role for BAT and white adipose tissue plasticity in wound healing and post-surgical energy expenditure. As fasting duration and sex represent important potential confounders in metabolic studies, it is important to note that the fasting times of the surgery model were shorter (2-4h) than for the acute/repeated acute models (10-12h) and include exclusively female mice. These differences limit direct comparability between the surgery and other stress models. Nevertheless, the magnitude of the observed metabolic differences is unlikely to be explained by these factors alone^44,45^. Importantly, our research indicates that surgery induces greater metabolic activation than anesthesia alone, corroborating previous findings that show increased adipose tissue activity during wound healing^46^. In addition to upregulation of classic thermogenic genes such as *Ucp1* and *Dio2*, we found that *Lpl* was remarkably upregulated in response to surgical trauma. This suggests a vital role for adipocyte-derived *Lpl* in BAT activity following injury^47^. We also observed post-surgical metabolic remodeling of iWAT and loss of volume, which has previously been associated with surgical stress^48^, reinforcing our findings. This might be associated with plasticity of iWAT as a potential energy and signaling fuel and browning for surrounding and distant BAT. To resolve the substrate utilization, we studied fatty acid metabolism using a long-chain fatty acid radiotracer, [^18^F]FTHA. We observed a substantially lower increase in uptake compared to the glucose derivative, [^18^F]FDG, indicating that glucose is the primary substrate supporting acute BAT activation. This is consistent with previous findings from Park et al.^49^, who reported glucose and lactate as the main energy sources for BAT activity, reflecting the rapid substrate demand for oxidative phosphorylation. Additionally, the increased energy demand of BAT may also be reflected by predominance of upregulated genes, consistent with the bioenergetic costs associated with increased transcriptional turnover^50^. Furthermore, by using a novel mitotyping approach^27,28^, we identified coordinated changes in mitochondrial gene and protein expression, characterized by increased OXPHOS and FAO following repeated cooling stress. These findings further highlight the mitochondrial remodeling as part of the adaptive response of adipose tissue to the increased energy demand of repeated stress exposure. In the surgery model, we observed a trend toward a prolonged metabolic shift toward increased fatty acid uptake, suggesting a transition to more energy-efficient substrate utilization during recovery or after stress exposure. Ucp1 expression correlated only during acute conditions with [^18^F]FDG uptake, whereas it correlated strongly with [^18^F]FTHA uptake, suggesting that Ucp1 is more relevant in adaptive processes than the rapid remodeling and substrate demand^51^. Alternative pathways, such as mTORC1^52^ and de novo lipogenesis^53^, likely contribute to this observation by redirecting glucose into biosynthetic processes. Conclusively, we present [^18^F]FTHA and [^18^F]FDG PET imaging as a robust imaging pair to assess adipose tissue remodeling and substrate utilization *in vivo*.

Beyond adipose tissue, we also observed changes in bone marrow and lymph node metabolism, indicating immune system involvement, which is known to be affected especially by repeated acute stress^54^. As immune cell function is closely linked to the metabolic state, we identified increased [^18^F]FDG uptake in splenic cell fractions, and reduced B-cell numbers using RadioFlow^55^. We also observed a trend towards decreased T-cell [^18^F]FDG uptake in bone marrow. Given the limited sample size, these findings should be considered exploratory and interpreted with appropriate caution. Nevertheless, they highlight the potential applicability of this novel method for investigating immunometabolism. Previous studies have also hypothesized post-surgical weight loss, possibly mediated by macrophages^56^, as BAT-associated macrophages have been shown to regulate sympathetic innervation and thermogenesis^57^.

Centrally, stress exposure reduced global brain glucose uptake by about 30% ^44,58^, while inducing stressor-specific regional alterations that intensified with severity. This is consistent with a redistribution of metabolic resources toward peripheral survival pathways and with imaging patterns linked to adverse outcomes, as recently described^2,59^. Additionally, the reduction in brain uptake might also be attributed to increased BGL levels in the early uptake phase or altered blood flow as an effect of stress^60^. This presents an approach that links neural [^18^F]FDG hyper- and hypometabolism to specific stress models. However, there are two main limitations in mice. First, the spatial resolution of mouse brain imaging, and secondly, the need for anesthesia, which we also found to be a crucial confounder of metabolic activity^61^. Nevertheless, for translational purposes, we performed our study entirely under fasting conditions and kept the protocol as close to the human clinical standard. With advances in total-body PET imaging in the clinical setting, we are confident that the preclinical establishment and brain analysis will drive future translational research to investigate brain activity in patients with cancer^5,26^ or other diseases^62^.

Next, we combined our imaging results at the total body level. Network-based analyses^23,24^ revealed stress-specific brain–adipose metabolic intra-organ connectivity, providing a framework for metabolic phenotyping of stress. Previous studies have shown the capability of network approaches in preclinical models, including neuroimaging^63,64^ or bone/skeletal metabolism^65,66^. Here, we move beyond single-organ analysis and present a brain-adipose tissue axis in acute and repeated acute stress models. However, network analysis, particularly in preclinical studies, is inherently constrained by sample size, which can affect the stability and robustness of the network estimates and therefore requires careful methodological consideration.

Taken together, our findings demonstrate that stress induces coordinated metabolic remodeling across adipose tissues and brain regions that can be quantified minimally invasively using PET imaging. By combining OMICS with total body PET, we establish BAT activation as a central and sensitive marker of the stress response, showing that glucose metabolism is the primary driver of acute activation that shifts to fatty acid substrate utilization and tissue remodeling upon repeated exposure or recovery. In addition, we reveal stressor-specific changes in regional brain metabolism and demonstrate that network-based approaches can depict these changes at the total body level. These results define a new role for PET imaging and imaging-based network analysis as tools for mechanistic and translational investigation of stress biology, providing a foundation for future studies aimed at improving diagnosis, monitoring, and intervention in stress-related diseases.

## Methods

### Animal experiments and ethics

All conducted experiments were ethically approved by the Austrian BMBWF (GZ 2021-0.030.763 and GZ 2023-0.062.018). As an animal experiment, this study followed the ARRIVE guidelines^67^ as best practice. Animals were randomly assigned to experimental groups (no specific randomization sequence used), and data were analyzed with the investigator blinded to group identity whenever possible. Cage location was kept constant throughout all experiments, and the order of treatments and measurements was mixed on each experimental day. Only the principal investigator was aware of the group allocation on the experimental days and during the outcome assessment. Male and female C57BL/6J-mice (N=90, 46m/44f) were supplied by Tierhaus Himberg (Medical University of Vienna) or Janvier (surgery) at 5-12 weeks old and allowed to acclimate at the animal facility of the Preclinical Imaging Lab for two weeks before any further experimental procedures were conducted. Due to logistical and experimental constraints, sex representation and age differ between the experimental models. Specifically, the surgery model included only female mice at 9 – 12 weeks old, whereas the acute and repeated acute stress models included both sexes and were 18 – 22 weeks old. Also, mice were fasted before imaging for 10-12 h for acute/repeated acute stress models or 2-4 h for the surgery model, as a shorter fasting period in the surgical cohort was implemented to minimize additional physiological stress during postoperative recovery. Mice were housed in groups of five, with some exceptions up to eight, separated by sex, in an IVC system (SmartFlow, Tecniplast GR900) and monitored as well as tail marked every other day. The housing conditions were maintained throughout the experiments with a temperature of 22 ± 2 °C, a light-dark cycle of 12/12 h, and *ad libitum* access to food, standard diet (Rod16-A, LASQCdiet®), and water. Cages were enriched with gnawing wood as well as plastic houses and tunnels. Body temperature was assessed either under isoflurane anesthesia using a rectal probe on the µPET/CT scanner bed or in fully conscious mice using an infrared thermometer (Sensian^TM^ 7, Braun). The IR thermometer was preferred for regular monitoring due to its non-invasiveness^68^. To ensure adequate precision, we measured the animals at least three times at the same site (abdominal area between the hind limbs) until the measurement was constant. The IR thermometer was cleaned between mice to prevent inaccuracy caused by contamination with urine and feces.

### Stress models

In a ventilated Styrofoam box, mice were individually placed in an ice-cooled cage at 4°C for 1 hour to induce acute cooling^69^. The mice were immobilized (acute restraint) by placing each mouse for one hour inside a modified 50mL Falcon® tube with several ventilation holes. The restrainers were placed on a heating pad maintained at 37 °C. In repeated acute stress models, the basic procedure was homologous to that in the acute model but was conducted once per day for one week. The repeatedly acutely stressed mice were treated like the control group on imaging day. Control animals were used to handling, and any additional distress was prevented whenever possible. To induce surgical stress, female C57BL6/J mice received 125 μg/g bodyweight Ketamine (Ketasol, Livisto) and 5 μg/g bodyweight Xylazine (Rompun®, Bayer) i.p. as anesthesia and were placed on a heating pad for the whole course of the surgery and wake-up phase. Vitamin A-containing eye cream (Vit-A-Vision®, OmniVision GmbH) was applied, and the mice received s.c. injection of 5 μL/g bodyweight of Meloxicam (Metacam®, Boehringer Ingelheim) for postoperative analgesia. Body hair was shaved with a sterile surgical blade (Swann-Morton), and Hardened Fine Scissors (Fine Science Tools) were used to cut an incision of 1 cm length on each flank. Subcutaneous connective tissue was cut with scissors, and subsequently, wounds were closed with a sterile Vicryl suture (6-0, BV-1 needle 30 cm violet) (Ethicon, Johnson & Johnson). Postoperative wound healing and behavior were assessed daily.

### Micro positron emission tomography/computed tomography imaging (µPET/CT) and magnetic resonance imaging (MRI)

[^18^F]FDG was produced in-house at the AKH Vienna with a GE Healthcare™ FASTlab Module using a cassette system, fulfilling the quality for human use according to the European Pharmacopoeia. [^18^F]FTHA was also produced in-house using a SOFIE Elixys Module^70^ with a radiochemical purity ≥ 97%. [^18^F]FDG and [^18^F]FTHA were diluted in a laminar air flow with sterile aqueous 0.9% NaCl solution to achieve an activity of 21 ± 3 MBq and 24 ± 4 MBq with an injection volume of 125 ± 25 µL and 142 ± 37 µL, respectively. Prior to imaging, the mice were fasted (see animal experiments and ethics paragraph for details), and their weight, length, and blood glucose levels (BGL) were recorded. For radiotracer injection, mice were anesthetized with Isoflurane (1-2% v/v in oxygen). Subsequently, [^18^F]FDG or [^18^F]FTHA was injected via the tail vein, and the radiotracer was allowed to distribute in awake status for 60 min. The control, repeated acute stress, and surgery groups were returned to their regular cages. In contrast, mice in the acute stress groups were directly exposed to the treatment after injection while awake. After the distribution phase, BGL was measured, and the mice were placed head-first on a heated bed (Medres Medical Research) with a fixed head position in the µPET/CT scanner (Inveon, Siemens^©^) for image acquisition covering the entire body within the field of view under continuous flow of Isoflurane. Static PET scans were acquired for 10 min, and dynamic images were acquired for 70 min. All images were attenuation-(CT-based), scatter-, decay-, and dead time-corrected. The detector was normalized during weekly quality control, in addition to best-practice daily quality controls. The reconstruction algorithms employed were OSEM3D/MAP for PET images and Feldkamp cone-beam for CT images. The final dimensions of the PET images were as follows: voxel size 0.388 × 0.388 × 0.796 mm (x/y/z) and image size 256 × 256 × 159 (x/y/z). The CT images had a voxel size of 0.0975 mm (x/y/z) and an image size of 1024 (x/y/z). PET/MR imaging data were acquired on a 9.4 T/30 MRI system (Bruker BioSpec) equipped with a PET insert (Si 198), using a homologous animal handling procedure for PET/CT acquisition. Anatomical MR images were obtained using a T□-weighted Turbo RARE sequence (coronal, fat-suppressed) with a phase-encoding direction along the row and four signal averages (echo time□=□34.2□ms, repetition time□=□5000□ms, and flip angle□=□180°). PET data were reconstructed using a 3D MLEM algorithm and were decay-, scatter-, randoms-, and dead-time–corrected and calibrated prior to analysis. The resulting voxel size was 0.137□×□0.640□×□0.251□mm (x/y/z) with an image size of 256□×□40□×□256 (x/y/z) for the MR images and a voxel size of 0.5mm (x/y/z) and an image size of 180□×□180□×□300 (x/y/z) for the PET images.

### Human PET/CT study

The study was conducted at the Division of Endocrinology and Metabolism, Department of Medicine III, Medical University of Vienna in accordance with the principles of the Declaration of Helsinki. The original human PET/CT study received approval from the Ethics Committee of the Medical University of Vienna (approval no. 1032/2013 and no. 1071/2017). Written informed consent was obtained from all participants before enrollment in the original study. 58 healthy individuals (n = 25 female) underwent pre- and post-cooling vest treatment [^18^F]FDG PET/CT imaging^71^. In short, the patients were fitted with a water-perfused cooling vest which covered the whole torso (CoolShirt Systems, Stockbridge, Georgia, USA). The temperature was gradually decreased until shivering was detected by electromyography (EMG Quattro, OT Bioeletronica, Torino, Italy) or the participant reported shivering and severe thermal discomfort, respectively. Volunteers received 2.5 MBq per kilogram of bodyweight of [^18^F]FDG intravenously followed by another 60 minutes of cold exposure during the tracer uptake^72^.

### Image analysis

PET images were quantified as standardized uptake value body weight (SUV_mean_) using PMOD software (PMOD Technologies LLC, Ver. 3.8). PET images were co-registered with the reference CT (reduction: 2/2/2; x/y/z) or MR image using automated rigid matching. Trilinear Interpolation was performed to match the voxel sizes of the CT image, resulting in a final voxel size of 0.195mm (x/y/z) and an image size of 512 (x/y/z). Interscapular and ventral spinal brown Adipose Tissue were delineated with an interscapular and ventral spinal spherical volume of interest (VOI) and subsequent application of the iso-contour tool to include regions with intensities greater than 70-80% of the maximum intensity within the VOI, based on the PET signal. Inguinal white adipose tissue (iWAT) was segmented by drawing contours in several planes, subsequently using an interpolation tool and iso-contouring of the CT signal to only include adipose tissue (<-100 HU). Inguinal lymph nodes were segmented in the iWAT using a sphere and iso-contouring to exclude any adipose tissue. Bone marrow was delineated by drawing a VOI plane-by-plane inside two vertebrae between the kidneys and the urinary bladder to prevent spillover. Muscle, liver, and spleen were segmented as representative spheres inside the quadriceps, left lateral lobe, and flank. The blood pool (inferior vena cava) was segmented by placing a sphere (0.5 mm radius) attentive to unwanted spillover affecting our dynamic data evaluation.

PET images were spatially normalized with rigid matching to the T2 mouse brain atlas^73^ (provided by PMOD) and masked to include only voxels within the atlas. Normalization was performed with proportional scaling^74^. To analyze the brain scans, we performed a voxel-wise comparison using statistical parametric mapping (SPM12) and a regional SUV_ratio_ quantification via the M. Mirrione Brain Atlas. We used a one-way ANOVA model and subsequent t-contrasts to compare the different stress models in SPM. Exploratory voxel-wise results were rendered with the PMOD 3D tool (Ver. 4.4), and full results are provided in the supplementary information. Regional brain atlas comparisons were corrected for multiple testing using Benjamini–Hochberg FDR (5% across all regions and contrasts within each analysis).

According to Lanz et al.^75^, we used an image-derived input function (IDIF) from the vena cava to evaluate our dynamic imaging data. The plasma-to-blood ratio was calculated with a previously published monoexponential correction function^76^. The parental fraction was assumed to be 1. Time-activity curves (TACs) were extracted as kBq/cc with Pmod. As a kinetic model, FDG Patlak was used to fit the Conc. (Tissue)/(Plasma) to AUC (Plasma)/Conc. (Plasma) from 5-35 min to limit potential early-frame inaccuracy (bolus, dispersion) and late-frame effects of FDG-6P dephosphorylation^77^. The goodness of the fit was assessed with R^2^ in Excel. Blood glucose levels, lumped constants (Brain: 0.6^77^, BAT: 1.14^78^), and Ki were used to calculate MR_glc_.

### [^18^F]FDG network analysis

The SUV_mean_ was used for comparison of brain and adipose tissue. A cut-off of r = 0.25 was pre-specified across cohorts as a cut-off. To justify it empirically, we benchmarked the threshold from r = 0 to r = 0.5 (step 0.05) and tracked preserved edges per cohort. Density and general connectivity declined steadily in all groups, with the biggest decrease between r = 0.10 and 0.30. Partial correlation networks of regional brain uptake using only significant (P < 0.05) connections were constructed using data filtered for stress-related regions. Standardized Uptake Values (SUVs) for analyzing brain regions were normalized to total brain uptake (SUV_ratio_). For all networks, further nonparametric bootstrapping of edge weights (1,000 resamples/cohort) was performed. Most confidence intervals were wide and overlapped zero, reflecting genuine estimation uncertainty at n = 12–20/group. All analyses were conducted in JASP (version 0.95.2). Networks were compared by correlating (Pearson correlation) their weight matrices^24^.

### *Ex-vivo* biodistribution

After imaging, mice were sacrificed by cervical dislocation. Complete dissection, as used for *ex vivo* biodistribution, included the following organs: heart, lung, liver, spleen, kidney, adrenal glands, pancreas, bladder, stomach, small intestine, large intestine, sexual organs, muscle, visceral WAT, inguinal WAT, iBAT, bone, thymus, thyroid, and brain, as well as blood and urine. We used an automated gamma counter (Hidex) to quantify radiotracer accumulation. The gamma counter was regularly calibrated with fluorine-18, and after decay correction, the results are expressed as the percentage of the injected radioactivity dose per gram of tissue (%ID/g).

### Transcriptomics

RNA extraction, purification, and sequencing were performed by the Genomic Core Facility at the CEITEC – Central European Institute of Technology (Masaryk University, Brno) using an Illumina NextSeq 500/550. The reads were mapped to the mouse genome (Ensembl GRCm39 release 111) with STAR aligner (version 2.5.2b)^79^ and subsequently sorted and indexed with Samtools (version 1.17)^80^. Deduplication via UMIs was performed with UMI Tools (version 1.1.4)^81^, and reads were counted using featureCounts (version 2.0.1)^82^. Data quality was assessed using FastQC (version 0.11.8) throughout the analysis. Low-abundance genes (counts < 5 in ≥ 2 samples) were removed, and differential gene expression analysis was performed in R (version 4.3.2) using DESeq2 (version 1.42.0)^83^ with BiomaRt (version 2.58.0) to map gene names to annotations and corrected for multiple testing via Benjamini-Hochberg. Genes were ranked according to Wald’s statistics, and gene set enrichment analysis was performed with fgsea^84^ using the Kyoto Encyclopedia of Genes and Genomes (KEGG) database gene set^85^ and clusterProfiler^86^. Results were displayed as volcano, enrichment, and bubble plots in R using enrichplot and ggplot2.

### Proteomics

Approximately 1-50 mg of tissue was combined with 50-150 µL 1x SDT buffer. Samples were disrupted by glass beads in a sonication device (Bioruptor^®^ Pico, Diagenode). For QC purposes, 1D-SDS-PAGE gels (12 %, CBB-G250 staining) were done, with a protein mixture of 5 µL loaded into each gel lane. Sample preparation was done by loading approximately 50 µg of protein onto FASP membranes (30 kDa cut-off, including DTT & Iodoacetamide, Incubation: Trypsin 18 h at 37 °C). The peptide mixture was purified using ethyl acetate extraction, resulting in a final solution of 90 µL. QC and semi-quantitative analyses of the peptide samples were done with LC-MS/MS on an RSLCnano + QTOF (Bruker) system. Quantitative LC-MS/MS analyses were done using nanoElute (Bruker) online connected to a timsTOF Pro system (Bruker) with a 104 min-long linear LC gradient. MS and MS/MS spectra were recorded in a time of flight (TOF) analyzer using data-independent acquisition (DIA) mode utilizing trapped ion mobility (TIMS) in the precursor m/z range of 400-1000. DIA LC-MS data processing was done using the DIA-NN application (Ver. 1.8). The library free search mode was used using the following protein databases: cRAP contaminant database (based on http://www.thegpm.org/crap/, 111 protein sequences in total, Ver. 1705) and UniProtKB_Mouse_can (taxonomy: Mus musculus, taxon ID: 10090, Ver. From 16.6.2021, 22,001 protein sequences in total). Match between runs (MBR) was used across the whole dataset. Database search results were set to follow FDR thresholds of 1% for precursor and protein group levels. DIA-NN was used to construct the protein groups list utilizing proteotypic and non-proteotypic peptides. MaxLFQ intensities were calculated based on identified peptides and extracted as normalized LFQ intensities (log2 transformed). Protein group IDs had to be present in at least 75% of all replicates to be used for further analysis. We used the LIMMA package^87^ in R (moderated t-test, Benjamini-Hochberg correction) with subsequent contrasts to calculate group differences. Proteins were mapped to Gene IDs using BiomaRt (version 2.58.0) and ranked by t-value for subsequent gene set enrichment analysis with fgsea^84^, using the KEGG database^85^ via clusterProfiler^86^. Results were shown as volcano and bubble plots using the ggplot2 package in R.

### Mitochondrial pathway and mitotyping analysis

Mitochondrial transcriptomic profiles were analyzed in R using an adapted implementation of the mitotyping framework described by Monzel et al.^27^. Mitochondrial genes and their functional pathway annotations were obtained from the mouse MitoCarta3.0 database^28^. MitoCarta3.0 assigns mitochondrial genes to 149 hierarchically organized MitoPathways covering major aspects of mitochondrial biology. Genes with fewer than 10 counts in at least three samples were excluded prior to downstream analysis. Following normalization of the transcriptomic data, MitoPathway scores were calculated for each sample as the mean normalized expression of all detected genes assigned to a given MitoPathway. Mitochondrial Pathway Priority Scores (mitoPPS) were subsequently calculated following the approach of Monzel et al. For each sample, pairwise ratios were calculated between all MitoPathway scores. Each pathway ratio was normalized to the mean ratio across all samples, generating a corrected pathway ratio. For each MitoPathway, corrected ratios relative to all other pathways were averaged to obtain the corresponding mitoPPS. Thus, mitoPPS represent the relative prioritization of individual mitochondrial pathways within the mitochondrial transcriptomic profile rather than absolute pathway expression. Principal component analysis (PCA) was performed on mitoPPS using the prcomp function in R after scaling pathway scores to unit variance. PCA loadings were used to identify mitochondrial pathways contributing most strongly to the major axes of variation. In addition to PCA across all experimental conditions, separate PCAs were performed for selected experimental groups to characterize the dominant mitochondrial variation within each comparison. PCA is unsupervised; therefore, subgroup-specific PC1 loadings describe pathways contributing to the principal axis of variation within the respective subset and were not interpreted as direct differential-expression effects relative to the control group. All analyses were performed in R version 4.6. Data processing and visualization were conducted using the tidyverse (v2.0.0)^88^, ggfortify (v0.4.19), ComplexHeatmap (v2.28.0), circlize (v0.4.18), and fs (v2.1.0) packages. DESeq2 (v1.52.0)^83^ was used for transcriptomic normalization and differential expression-related preprocessing.

### Quantitative real-time PCR (qRT-PCR)

Total RNA was extracted from snap-frozen tissues (10-50 mg) with TRIzol™ (Invitrogen, ThermoFisher) and RNeasy Lipid Tissue Mini Kit (Qiagen). Samples were homogenized with a rotor-stator homogenizer (Ultra-Turrax T18, IKA) at 19000rpm for 40 seconds and with a Precellys Homogenizer for 2 × 15sec at 5500rpm/min (Bertin Technologies). RNA concentration and purity were assessed with a multimode plate reader (Infinite® 200 Pro, Tecan) and a Spectrophotometer (NanoDrop™ 2000, ThermoFisher). cDNA Synthesis of 1000ng of RNA was performed with the iScript™ Advanced cDNA Synthesis Kit (Bio-Rad) and high-capacity cDNA reverse transcriptase kit (Applied Biosystems). Afterward, cDNA samples were diluted 1:25 with RNase-free water (Invitrogen, Thermo Fisher). Real-time PCR was performed with a CFX Connect™ Real-time PCR Detection System (Bio-Rad) using PrimePCR^TM^ Assays (Bio-Rad, *Acadl, Acadm, Acadvl, Adrb2, Adrb3, Cd36, Crebbp, Dio2, Fabp4, Lpl, Mt2, Pdk4, Pfkfb1, Pnpla2, Ppara, Pparg, Ppargc1a, Prdm16, Sgk1, Slc2a1, Tbp, Ucp1,* for further details see 10.5281/zenodo.21889568) with SsoAdvanced Sybr Green Supermix (Bio-Rad) in 96-well plates. All samples were run in technical duplicates, with NTCs for each primer pair, and two housekeeping genes per plate to account for interplate variability. Cq values were extracted using CFX Manager (Ver. 3.1, Bio-Rad) and evaluated by the 2^-ΔΔCt^ method in Excel, with Tbp as the reference gene to obtain relative expression changes.

### Enzyme immunoassays

Blood samples were collected at multiple time points. Baseline levels were measured between 9 and 10 am on randomly selected days, with at least two weeks separating sampling and imaging. Within the imaging procedure, blood sampling was performed during the application, after stress exposure (via tail vein), and after the scan (via cardiac puncture). Serum was collected, frozen, and stored at -80 °C for subsequent analyses. According to the manufacturer’s manual, a Corticosterone ELISA kit (ADI-900-097, Enzo Life Sciences) was used to quantify corticosterone levels in serum samples. The results were obtained using a plate reader (Biotek® Synergy HTX, Agilent) with a dual-wavelength method, with readouts at 405 nm (signal) and 580 nm (background), each blanked against the corresponding blank wells and then subtracted from one another. A 4-parameter logistic curve was fitted to the standards and used to calculate sample concentration using integrated software (Biotek Gen5, Agilent).

Feces samples were collected in the morning and evening over one week to account for repeated acute stress and circadian rhythm effects. Besides pooled feces collection in the standard IVCs, individual sampling was done in specifically modified cages^89^. Corticosterone metabolites were determined in feces samples with a 5α-pregnane-3β, 11β, 21-triol-20-one EIA following methanol extraction (see Touma et al.^43^ for details) at the Department of Biological Sciences and Pathobiology (Experimental Endocrinology) of the Veterinary University of Vienna.

### Histology

iBAT and iWAT tissues were cut into small cubes (3 – 5 mm) and pretreated with 15% and 30% sucrose overnight. Then tissues were embedded in OCT compound and cut with a cryostat (Crystar NX70, Epredia) on coated slides (SuperFrost Plus, Epredia) into 8-10 µm sections. We used three technical replicates per slide and fixed the slides for 10 minutes in ice-cold Ethanol (96% v/v). H&E staining was performed according to a standard protocol, using Gill’s III Hematoxylin (Merck) and Eosin-Y solution (0.5% aqueous, Merck), and mounted with VectaMount® (VectaLabs).

The iBAT and iWAT sections were fixed in cold 4% PFA (Roti Histofix, Carl Roth) for 10 minutes, and peroxidase was blocked by incubation in 3% H_2_O_2_ (30% H_2_O_2_, Merck) for 10 minutes. We blocked the sections with Avidin, Biotin (Blocking Kit, VectaLabs), BLOXALL™ (VectaLabs), and 5% BSA (Sigma-Aldrich) in PBST. Additionally, we permeabilized the sections with 0.3% Triton X-100 in PBS. Primary UCP-1 Antibody (Abcam, EPR20381, ab209483) was applied overnight at 4°C in a 1:20000 dilution in PBST. The secondary Antibody (biotinylated horse anti-mouse/rabbit IgG, VECTASTAIN Elite ABC-HRP Kit, Vector Laboratories, PK-6200) was used at a dilution of 1:500 for 2 hours at room temperature. Slides were treated with AB mixture (Vectastain ABC Kit, VectaLabs), and DAB substrate (ab64238, Abcam) was applied for 3 minutes. Counterstaining was performed with Gills III Hematoxylin (Diluted 1:1 with dist. H2O). Slides were dehydrated in ethanol and n-butyl acetate, then mounted in VectaMount. Analysis of all sections was performed using an Olympus VS200 slide scanner with the default IHC brightfield 20x protocol. Images were post-processed and analyzed with OlyVIA Software (Ver. 4.2) and QuPath (Ver. 0.6.0). After automated estimation of stain vectors, DAB-positive area was quantified (DAB ≥ 0.15 OD) and compared to total tissue area (Average channels ≤ 200 – 220 OD).

### Fluorescence activated cell sorting (RadioFlow)

Cell suspensions of the spleen were prepared by mechanical straining through a 70 µm and 40 µm Falcon® cell strainer. Bone marrow was flushed out from the femur and tibia in FACS buffer containing 1% FBS and EDTA through a strainer, and for both tissues, lysis of erythrocytes was conducted. Aliquots were measured for radioactivity concentration and total cell number before staining. Cells were pre-treated for Fc-blocking (CD16/32, 1:200, eBioscience 14-0161-85) and labelled ex vivo with the following fluorescent monoclonal antibodies for spleen: CD11b (BV605, 1:800, Biolegend 101237), CD4 (FITC, 1:400, eBioscience 11-0041-82), B220 (PE-Cy7, 1:400, eBioscience 25-0452-82), CD19 (PE, 1:400, BD 557399), CD3 (APC-Cy7, 1:200, Biolegend 100222), CD8 (APC, 1:400, eBioscience 17-0081-82) and bone marrow: CD11b (BV605, 1:800, Biolegend 101237), CD23 (eFluor450, 1:400, eBioscience 48-0232-82), CD3 (FITC, 1:400, BD 553061), B220 (PE-Cy7, 1:400, eBioscience 25-0452-82), Ly6G(PE, 1:2000, Biolegend 127608), CD117 (APCeFluor780, 1:200, eBioscience 47-1171-82), Ter-119 (APC, 1:400, Biolegend 116211). The samples were analyzed using a FACSAria™ Fusion Flow Cytometer (Becton Dickinson, USA), and the fluorescence data were analyzed with FlowJo (FlowJo, LLC, OR, USA). The radioactivity of the sorted cell fractions was measured in a gamma counter (Wizard2, PerkinElmer Inc., Waltham, MA, USA). Counts per minute (CPM) were expressed as a percentage of total CPM per 100,000 cells, next to the total number of cells in each population. The results of single-cell fractions are illustrated relative to the total counts of all cell fractions. The total number of cells in the sorted population was calculated within the cell sorting system.

### Statistics and Reproducibility

Statistical analysis and graphical visualization were performed with GraphPad Prism (Ver. 10.1.0). We employed regular two-tailed t-tests for statistical comparison of two groups and one-way or two-way analysis of variance (ANOVA) for comparison of three or more groups, with post-hoc Dunnett and Tukey tests to adjust for multiple comparisons. Paired t-test or repeated-measures ANOVA were used whenever applicable. Benjamini-Hochberg false discovery rate (FDR) correction was applied where a high number of multiple comparisons were performed in parallel. A p-value of < 0.05 was deemed statistically significant, and data are presented as mean ± SEM. Measurements were obtained from individual mice without pooling samples, and repeated measurements within mice were accounted for where applicable. No outlier criteria were set a priori, but two animals (ID 61, 62) were excluded from imaging and blood glucose analyses due to non-fasting conditions and two more (ID 91, 93) due to technical errors. Reproducibility of results was assured by randomization of experimental group allocation, blinding of the analyst (mouse ID only), and repetition of experiments across several months. Sample size was determined from the ethical approval with no explicit *a priori* sample size calculation performed. Due to technical and experimental feasibility, some experiments are underpowered, but this is explicitly mentioned when applicable with regards to the exploratory character of this study. Details regarding statistical testing are provided within the figure legends.

## Additional Information

### Data Availability

The mass spectrometry proteomics data have been deposited in the ProteomeXchange Consortium via the PRIDE^90^ partner repository with the dataset identifier PXD082804. The RNA-Seq data for this study have been deposited in the European Nucleotide Archive (ENA) at EMBL-EBI under accession number PRJEB124202. All source data for each displayed figure and additional raw data have been uploaded to a general repository (doi: 10.5281/zenodo.21889568)91. Any other raw data files, such as DICOMs, are available upon request from the corresponding author, but were not shared publicly due to file size and the lack of a suitable repository. Human data used in this study were previously published and are cited in the manuscript.

### Code Availability

Relevant code for the RNASeq, proteomics, and Mitotyping analyses is available via the general repository (doi: 10.5281/zenodo.21889568)91.

## Acknowledgements

Thank you to Stefanie Ponti, Anna Zacher, Lara Breyer, Johann Stanek, and Yago Lutz for their technical assistance with the animal experiments and data, as well as to the animal care staff and the Center for Biomedical Research and Translational Surgery at the Medical University of Vienna. We also thank the radiotechnology and radiochemistry team of the Division of Nuclear Medicine for [^18^F]FDG synthesis and the technical support and radiation protection management provided by Thomas Zenz and Andreas Krcal. Further thanks to Katherina Tillmann for providing custom-made mouse cages suitable for individual fecal sampling. Additional thanks to the Institute of Medical Statistics of the Medical University of Vienna for providing valuable statistical feedback. We acknowledge the CF Genomics and the CF Bioinformatics, supported by the NCMG research infrastructure (LM2023067 funded by MEYS CR), and the CF Prot of CIISB, Instruct-CZ Centre, supported by MEYS CR (LM2023042) and European Regional Development Fund-Project „Innovation of Czech Infrastructure for Integrative Structural Biology“ (No. CZ.02.01.01/00/23_015/0008175), for their support in obtaining scientific Transcriptomics and Proteomics data presented in this article. We also acknowledge the support of the Research Platform Medical Imaging (RPMI) of the Medical University of Vienna.

## Author Contributions

Conceptualization: MK, CV, MH; Methodology: MK, CV; TWA, RP, CK, UU, OK, FK, TWE, TH, JF,CM; Software: /; Validation: MK, JLW, PE, SMM; Formal Analysis: MK, CV, SMM, HV, ÖÖ, OK, CP, AG, CM; Investigation: MK, JLW, CV; Resources: MH, TWA, CK, RP, TH, FK, CV, CP; Data Curation: MK, CV, BG; Writing – Original Draft Preparation: MK, CV; Writing – Review & Editing: SG, MH, TWE, RP; CM; Visualization: MK, CV, BG, AG, CM; Supervision: CV, MH; Project Administration: SG; Funding Acquisition: MH, CV;

## Inclusion and Ethics

We support inclusive, diverse, and equitable conduct of research.

