## Supplementary_Information for "Stress-induced Metabolic Remodeling of Adipose and Brain Tissue Revealed by Positron Emission Tomography"

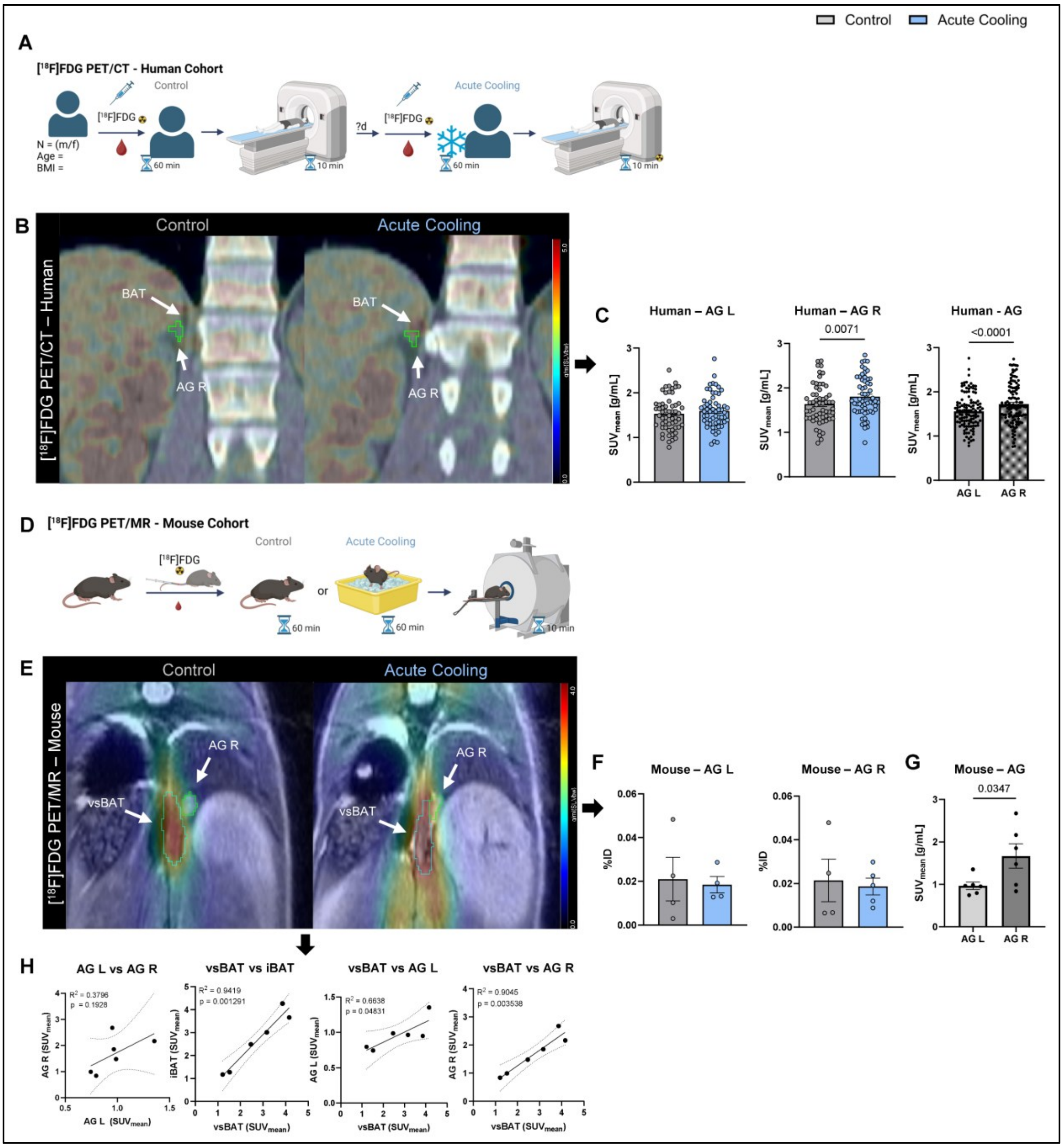


**Supplementary Figure 1: BAT depots and not adrenal glands show increased [18F]FDG uptake upon acute cold exposure in humans and mice**

**A.** Illustration of the clinical study design of acute cooling exposure of healthy individuals. The presented human data were previously published and are cited in the methods section of the main text (see Human PET/CT study). Created in BioRender. Krisch, M. (2026) https://BioRender.com/4972jxa.

**B**. Representative [18F]FDG PET/CT images from control scans (left) and after acute cooling exposure (right) in the human study cohort showing adrenal glands and the proximal BAT depots.

**C**. Quantification ($SUV_{mean}$) of [$^{18}$F]FDG uptake in adrenal glands in humans in control scans and after acute cooling exposure (n = 58).

**D**. Illustration of the preclinical imaging study for control and acute cooling intervention in C57BL/6J mice. Created in BioRender. Krisch, M. (2026) https://BioRender.com/4972jxa.

**E**. Representative [$^{18}$F]FDG PET/MR images in control (left) and acute cooling (right) mice showing adrenal glands and the proximal BAT depots of mice.

**F**. Quantification (% ID) of [$^{18}$F]FDG uptake in adrenal glands via ex vivo biodistribution of control (n = 4) and acute cooling (n = 4, AG L; n = 5, AG R) mice.

**G**. Quantification ($SUV_{mean}$) of [$^{18}$F]FDG uptake in adrenal glands via PET/MR analyses of mixed control and acute stress mice (n = 6).

**H**. Linear regression analyses of vsBAT, iBAT, and adrenal gland [$^{18}$F]FDG uptake in mixed control and acute stress mice (n = 6).

Data were analyzed using paired two-sided t-tests (**C, G**) and unpaired two-sided t-tests (**F**). Data are presented as individual values and as mean ± SEM, with p-values indicated as numbers above the bars.

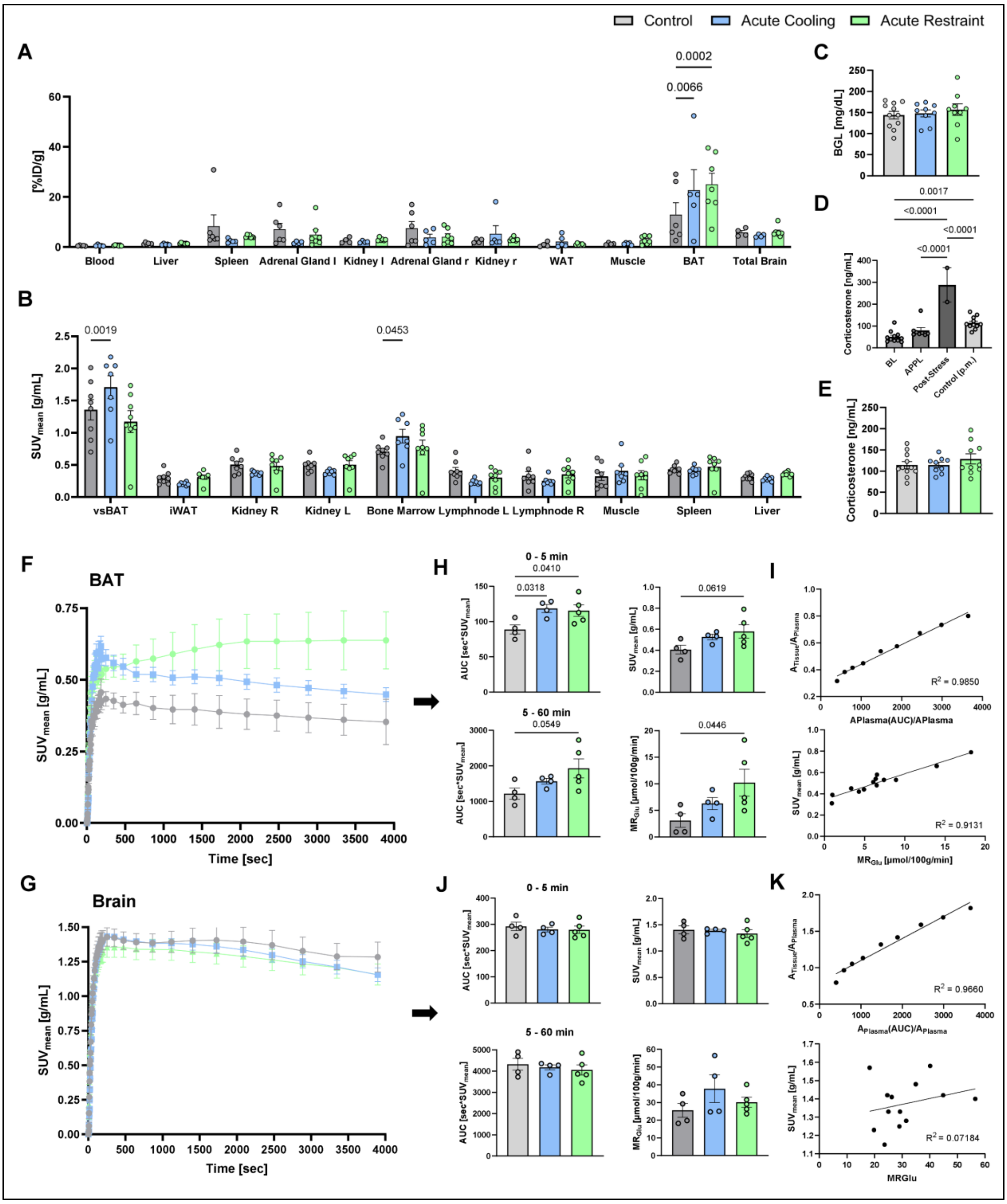


**Supplementary Figure 2: Dynamic [$^{18}$F]FDG µPET/CT reveals kinetics of iBAT and brain uptake in acute stress models.**

**A**. Quantification (%ID/g) of [$^{18}$F]FDG uptake via ex vivo biodistribution of control (n = 4-6), acute cooling (n = 4-5), and acute restraint (n = 6-7) mice.

**B**. Quantification (SUV$_{mean}$) of [$^{18}$F]FDG uptake via static µPET/CT imaging in multiple organs of control (n = 8), acute cooling (n = 7), and acute restraint (n = 8) mice.

**C**. Blood glucose levels (mg/dL) of control (n = 11), acute cooling (n = 9), and acute restraint (n = 9) mice.

**D**. ELISA analysis of serum corticosterone levels measured at baseline (n = 11), during radiotracer application (n = 8), after stress exposure (n = 2), and post-mortem (n = 11, control group only).

**E**. ELISA analysis of post-mortem serum corticosterone levels in control (n = 11), acute cooling (n = 9), and acute restraint (n = 10) mice.

**F, G**. Time-activity curves (TACs) of dynamic [$^{18}$F]FDG µPET/CT of iBAT (**F**) and brain (**G**) in control (n = 4), acute cooling (n = 4), and acute restraint (n = 5) mice.

**H, J**. Quantification of [$^{18}$F]FDG uptake via dynamic µPET/CT of iBAT (**H**) and brain (**J**) in control (n = 4), acute cooling (n = 4), and acute restraint (n = 5) mice as area under the curve in early (left, top) and late time frames (left, bottom), average $SUV_{mean}$ (right, top), and metabolic rate ($MR_{Glu}$) (right, bottom).

**I, K**. Representative Patlak plots (top) and linear regression of $SUV_{mean}$ and $MR_{Glu}$ (bottom) of iBAT (**I**) and brain (**K**) tissue. Regression analyses include control (n = 4), acute cooling (n = 4), and acute restraint (n = 5) mice.

Data were analyzed using two-way ANOVA (**A, B**) with Dunnett's post-hoc test, one-way ANOVA (**C, E, H, J**) with Dunnett's post-hoc test, and one-way ANOVA (**D**) with Tukey's post-hoc test. Data are presented as individual values and as mean ± SEM, with p-values indicated as numbers above the bars.

Control | Acute Cooling | Acute Restraint

**A**

| set p | set c | cluster p(FWE-corr) | cluster p(FDR-corr) | cluster equivk | cluster p(unc) | peak p(FWE-corr) | peak p(FDR-corr) | peak T | peak equivZ | peak p(unc) | x,y,z {mm} | x,y,z {mm} | x,y,z {mm} |
|---|---|---|---|---|---|---|---|---|---|---|---|---|---|
| **Hypermetabolism - Control vs Acute Cooling** | | | | | | | | | | | | | |
| 1.83E-01 | 3 | 9.20E-06 | 1.88E-05 | 2580 | 6.28E-06 | 6.09E-02 | 6.43E-01 | 5.40 | 4.19 | 1.38E-05 | -1.8 | 7.2 | -6.2 |
| | | 7.13E-01 | 8.76E-01 | 4 | 8.52E-01 | 6.73E-01 | 8.99E-01 | 3.72 | 3.21 | 6.74E-04 | 3.0 | 6.0 | -5.6 |
| | | 7.23E-01 | 8.76E-01 | 3 | 8.76E-01 | 6.76E-01 | 8.99E-01 | 3.72 | 3.20 | 6.81E-04 | 0.4 | 6.4 | -2.6 |
| **Hypometabolism - Control vs Acute Cooling** | | | | | | | | | | | | | |
| **set p** | **set c** | **cluster p(FWE-corr)** | **cluster p(FDR-corr)** | **cluster equivk** | **cluster p(unc)** | **peak p(FWE-corr)** | **peak p(FDR-corr)** | **peak T** | **peak equivZ** | **peak p(unc)** | **x,y,z {mm}** | **x,y,z {mm}** | **x,y,z {mm}** |
| 6.14E-02 | 4 | 4.38E-01 | 8.76E-01 | 56 | 3.94E-01 | 2.94E-01 | 8.98E-01 | 4.42 | 3.65 | 1.33E-04 | -2.4 | 5.0 | -2.0 |
| | | 6.37E-01 | 8.76E-01 | 14 | 6.91E-01 | 5.76E-01 | 8.98E-01 | 3.88 | 3.31 | 4.63E-04 | 2.6 | 4.8 | -1.4 |
| | | 6.64E-01 | 8.76E-01 | 10 | 7.44E-01 | 6.75E-01 | 8.98E-01 | 3.72 | 3.20 | 6.79E-04 | 1.8 | 4.6 | -0.6 |
| | | 7.23E-01 | 8.76E-01 | 3 | 8.76E-01 | 7.32E-01 | 8.98E-01 | 3.62 | 3.14 | 8.53E-04 | 2.6 | 3.8 | -1.0 |
| **Hypermetabolism - Control vs Acute Restraint** | | | | | | | | | | | | | |
| **set p** | **set c** | **cluster p(FWE-corr)** | **cluster p(FDR-corr)** | **cluster equivk** | **cluster p(unc)** | **peak p(FWE-corr)** | **peak p(FDR-corr)** | **peak T** | **peak equivZ** | **peak p(unc)** | **x,y,z {mm}** | **x,y,z {mm}** | **x,y,z {mm}** |
| 3.99E-03 | 6 | 1.01E-01 | 4.37E-01 | 264 | 7.28E-02 | 2.03E-01 | 4.91E-01 | 4.67 | 3.79 | 7.43E-05 | 0.6 | 3.2 | 4.6 |
| | | 3.13E-01 | 6.62E-01 | 99 | 2.56E-01 | 2.49E-01 | 4.91E-01 | 4.53 | 3.71 | 1.02E-04 | -3.2 | 4.8 | -3.8 |
| | | 4.53E-01 | 6.62E-01 | 52 | 4.12E-01 | 2.92E-01 | 4.91E-01 | 4.42 | 3.65 | 1.31E-04 | -0.4 | 5.0 | -0.4 |
| | | 4.76E-01 | 6.62E-01 | 46 | 4.42E-01 | 5.03E-01 | 6.46E-01 | 4.01 | 3.39 | 3.47E-04 | 3.2 | 4.6 | -4.0 |
| | | 5.94E-01 | 7.39E-01 | 21 | 6.16E-01 | 5.40E-01 | 6.46E-01 | 3.94 | 3.35 | 4.00E-04 | -1.6 | 3.0 | 5.2 |
| | | 6.95E-01 | 8.10E-01 | 6 | 8.10E-01 | 5.63E-01 | 6.46E-01 | 3.90 | 3.33 | 4.39E-04 | -0.8 | 5.4 | -1.6 |
| **Hypometabolism - Control vs Acute Restraint** | | | | | | | | | | | | | |
| **set p** | **set c** | **cluster p(FWE-corr)** | **cluster p(FDR-corr)** | **cluster equivk** | **cluster p(unc)** | **peak p(FWE-corr)** | **peak p(FDR-corr)** | **peak T** | **peak equivZ** | **peak p(unc)** | **x,y,z {mm}** | **x,y,z {mm}** | **x,y,z {mm}** |
| 6.14E-02 | 4 | 5.89E-01 | 6.07E-01 | 22 | 6.07E-01 | 5.93E-01 | 8.31E-01 | 3.85 | 3.29 | 4.94E-04 | 4.0 | 1.6 | -2.4 |
| | | 8.35E-02 | 2.38E-01 | 295 | 5.95E-02 | 6.10E-01 | 8.31E-01 | 3.83 | 3.28 | 5.27E-04 | 3.6 | 5.8 | -1.0 |
| | | 3.95E-01 | 6.07E-01 | 69 | 3.43E-01 | 6.28E-01 | 8.31E-01 | 3.80 | 3.26 | 5.66E-04 | -3.8 | 5.4 | -1.0 |
| | | 5.67E-01 | 6.07E-01 | 26 | 5.72E-01 | 7.04E-01 | 8.31E-01 | 3.67 | 3.17 | 7.62E-04 | -2.8 | 5.6 | 0.4 |
| **Hypermetabolism - Acute Cooling vs Acute Restraint** | | | | | | | | | | | | | |
| **set p** | **set c** | **cluster p(FWE-corr)** | **cluster p(FDR-corr)** | **cluster equivk** | **cluster p(unc)** | **peak p(FWE-corr)** | **peak p(FDR-corr)** | **peak T** | **peak equivZ** | **peak p(unc)** | **x,y,z {mm}** | **x,y,z {mm}** | **x,y,z {mm}** |
| 3.99E-03 | 6 | 2.75E-11 | 1.13E-10 | 7639 | 1.88E-11 | 5.96E-03 | 1.08E-01 | 6.76 | 4.82 | 7.03E-07 | -0.8 | 4.6 | 0.0 |
| | | 4.46E-01 | 9.38E-01 | 54 | 4.03E-01 | 5.28E-01 | 6.91E-01 | 3.96 | 3.36 | 3.83E-04 | 1.0 | 4.4 | -6.4 |
| | | 7.13E-01 | 9.38E-01 | 4 | 8.52E-01 | 7.30E-01 | 9.56E-01 | 3.62 | 3.14 | 8.48E-04 | 0.8 | 4.6 | -7.2 |
| | | 7.34E-01 | 9.38E-01 | 2 | 9.04E-01 | 7.35E-01 | 9.56E-01 | 3.61 | 3.13 | 8.66E-04 | -0.2 | 4.2 | -6.0 |
| | | 7.47E-01 | 9.38E-01 | 1 | 9.38E-01 | 7.48E-01 | 9.65E-01 | 3.59 | 3.12 | 9.13E-04 | -1.4 | 2.6 | -6.0 |
| | | 7.47E-01 | 9.38E-01 | 1 | 9.38E-01 | 7.63E-01 | 9.81E-01 | 3.56 | 3.10 | 9.72E-04 | -1.2 | 5.2 | -7.4 |
| **Hypometabolism - Acute Cooling vs Acute Restraint** | | | | | | | | | | | | | |
| **set p** | **set c** | **cluster p(FWE-corr)** | **cluster p(FDR-corr)** | **cluster equivk** | **cluster p(unc)** | **peak p(FWE-corr)** | **peak p(FDR-corr)** | **peak T** | **peak equivZ** | **peak p(unc)** | **x,y,z {mm}** | **x,y,z {mm}** | **x,y,z {mm}** |
| 1.70E-02 | 5 | 2.83E-11 | 9.65E-11 | 7626 | 1.93E-11 | 5.97E-03 | 1.03E-01 | 6.76 | 4.82 | 7.06E-07 | -1.6 | 7.2 | -6.2 |
| | | | | | | 5.79E-02 | 2.79E-01 | 5.43 | 4.21 | 1.29E-05 | -4.6 | 1.0 | -1.6 |
| | | 1.61E-02 | 1.84E-02 | 595 | 1.11E-02 | 2.41E-01 | 5.51E-01 | 4.55 | 3.73 | 9.70E-05 | 4.2 | 4.2 | -6.6 |
| | | 3.82E-04 | 6.52E-04 | 1475 | 2.61E-04 | 2.71E-01 | 5.51E-01 | 4.47 | 3.68 | 1.16E-04 | 5.0 | 3.4 | -1.6 |
| | | 2.62E-01 | 2.59E-01 | 123 | 2.07E-01 | 6.11E-01 | 8.07E-01 | 3.83 | 3.27 | 5.29E-04 | 3.0 | 5.8 | 0.8 |
| | | 7.47E-01 | 9.38E-01 | 1 | 9.38E-01 | 7.60E-01 | 9.74E-01 | 3.57 | 3.10 | 9.62E-04 | -1.2 | 6.4 | -2.4 |

**B**

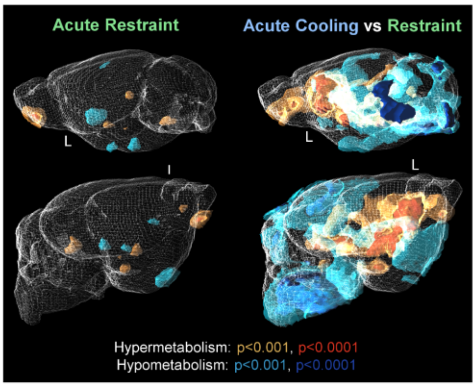


**Supplementary Figure 3: Voxel-wise brain analysis of acute stress models**

**A.** Summarized quantitative results of voxel-wise analysis performed with statistical parametric mapping (SPM12) using a one-way ANOVA model with post-hoc t-contrasts. Corresponding images are shown in Fig. 1F and Supplementary Fig. 3B.

**B.** Voxel-wise exploratory comparison of normalized brain [$^{18}$F]FDG uptake showing differences between control (n = 8) and acute restraint (n = 8) mice on the left and between acute cooling (n = 7) and acute restraint (n = 8) mice on the right, with thresholds of p < 0.001 and p < 0.0001 (uncorr.) to render significantly changed areas with the PMOD 3D tool (Ver. 4.4). The yellow and red areas indicate hypermetabolism, while the blue and dark blue areas indicate hypometabolism.

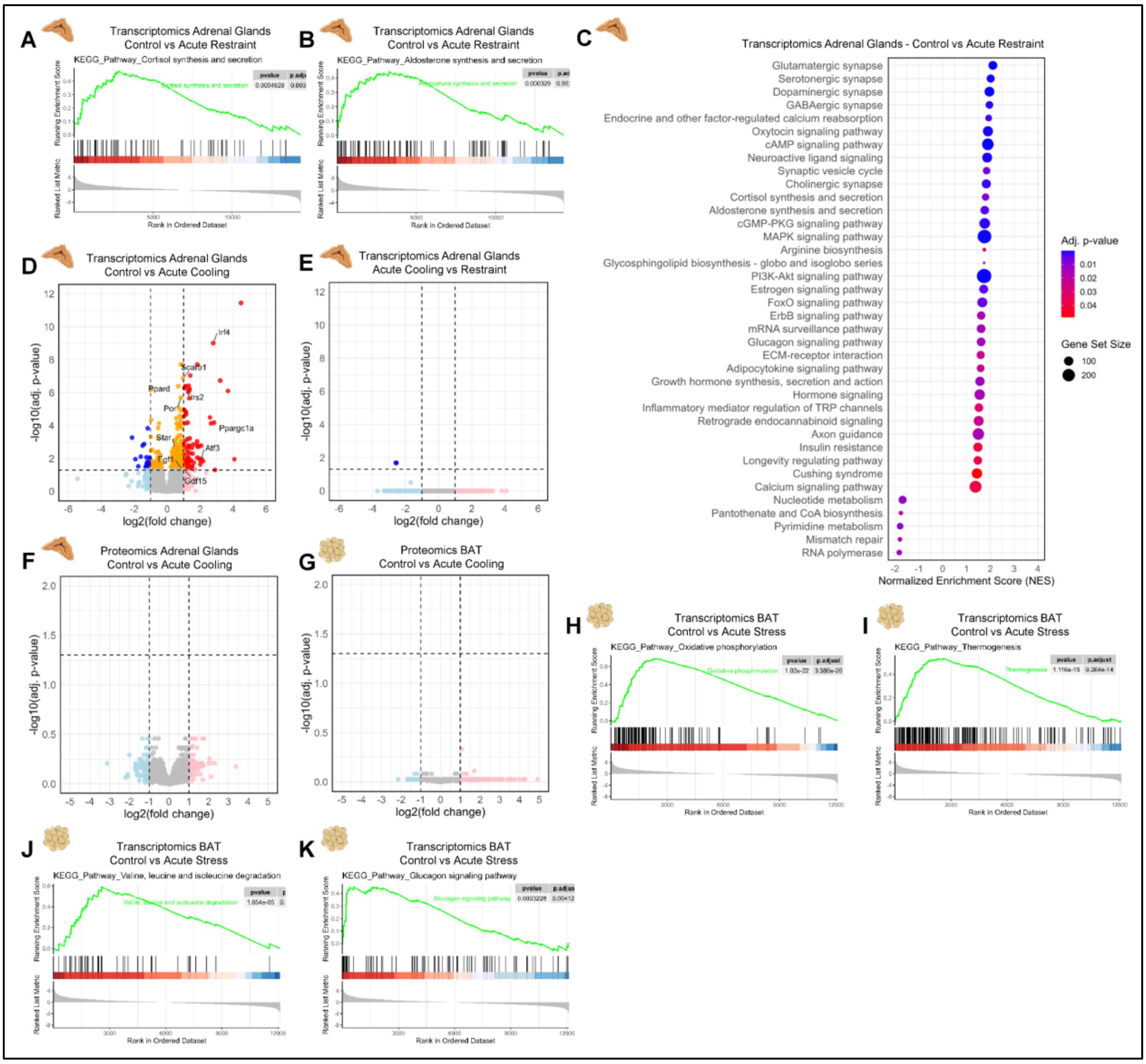


**Supplementary Figure 4: Bulk RNA-Seq and Proteomics analysis of Adrenal Glands and iBAT in acute stress models**

**A, B**. Enrichment plots of Cortisol (**A**) and Aldosterone (**B**) synthesis and secretion KEGG pathways in control (n = 5) vs acute restraint (n = 3) mice of adrenal glands via bulk RNA-Seq analysis. The headings in the Omics datasets indicate the direction of the comparison (A vs. B), with A as the control condition.

**C**. KEGG pathway gene set enrichment analysis of bulk RNA-Sequencing results of adrenal glands of control (n = 5) and acute restraint (n = 3) mice illustrated as a bubble plot showing normalized enrichment score (NES), adjusted p-value (adj. p-value), and gene set size.

**D, E**. Bulk RNA-sequencing results of adrenal glands in control (n = 5) vs acute cooling (n = 3) (**D**) and acute cooling (n = 3) vs acute restraint (n = 3) (**E**) mice. Differential gene expression was analyzed using DESeq2 with two-sided Wald tests and Benjamini-Hochberg correction and illustrated as a volcano plot (downregulated: blue, downregulated (ns): light blue, upregulated: red, upregulated (ns): light red, |log2FC| ≤ 1 & p-adj. ≤ 0.05: orange, no change: grey).

**F**. Proteomics results of adrenal glands in control (n = 5) and acute cooling (n = 3) mice, analyzed using LIMMA with two-sided moderated t-tests and Benjamini-Hochberg correction, illustrated as a volcano plot (legend see **D, E**).

**G**. Proteomics results of iBAT in control (n = 3) and acute stress (n = 3; cooling and restraint) mice, analyzed using LIMMA with two-sided moderated t-tests and Benjamini-Hochberg correction, illustrated as a volcano plot (legend see **D, E**).

**H-K**. Enrichment plots of Oxidative Phosphorylation (**H**), Thermogenesis (**I**), Valine, Leucine, and Isoleucine degradation (**J**), and Glucagon Signaling (**K**) KEGG pathways in control (n = 3) vs acute stress (n = 3; cooling and restraint) mice of iBAT via bulk RNA-Seq analysis.

Pictograms of organs in A-K were taken from BioRender. Vraka, C. (2026) https://BioRender.com/d1uqx40.

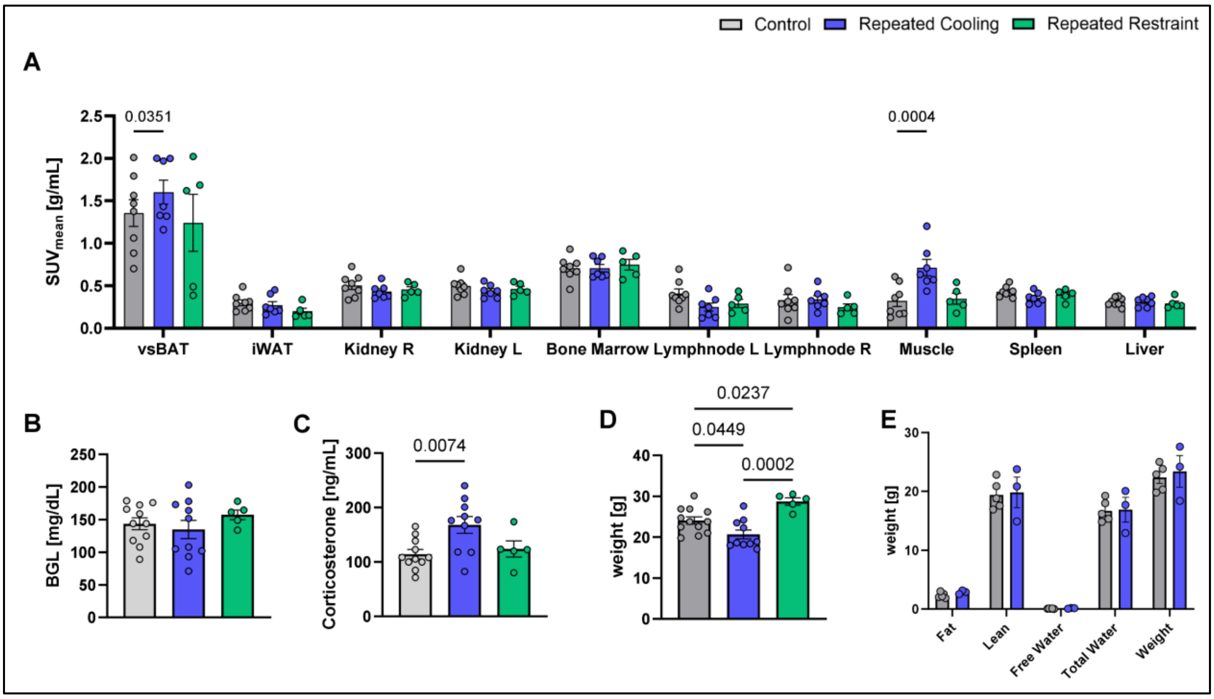


**Supplementary Figure 5: Imaging and health parameters of repeated cooling mice reveal increased vsBAT activity, corticosterone levels, and reduced bodyweight**.

**A**. Quantification ($SUV_{mean}$) of [$^{18}$F]FDG uptake via static µPET/CT imaging in multiple organs of control (n = 8), repeated cooling (n = 7), and repeated restraint (n = 5) mice.

**B**. Blood glucose levels (mg/dL) in control (n = 11), repeated cooling (n = 10), and repeated restraint (n = 5) mice.

**C**. ELISA analysis of post-mortem serum corticosterone levels in control (n = 11), repeated cooling (n = 10), and repeated restraint (n = 5) mice.

**D**. Bodyweight (g) in control (n = 12), repeated cooling (n = 10), and repeated restraint (n = 5) mice.

**E**. Body composition analysis using Echo-MRI in control (n = 5) and repeated cooling (n = 3) mice.

Data were analyzed using two-way ANOVA (**A**) with Dunnett's post-hoc test, one-way ANOVA (**B, D**) with Tukey's post-hoc test, one-way ANOVA (**C**) with Dunnett's post-hoc test, and two-way ANOVA (**E**) with Šídák's post-hoc test. Data are presented as individual values and as mean ± SEM, with p-values indicated as numbers above the bars.

Control | Repeated Cooling | Repeated Restraint

**A**

| set p | set c | cluster p(FWE-corr) | cluster p(FDR-corr) | cluster equivk | cluster p(unc) | peak p(FWE-corr) | peak p(FDR-corr) | peak T | peak equivZ | peak p(unc) | x,y,z {mm} | x,y,z {mm} | x,y,z {mm} |
|---|---|---|---|---|---|---|---|---|---|---|---|---|---|
| **Hypermetabolism - Control vs Repeated Cooling** | | | | | | | | | | | | | |
| 3.08E-01 | 3 | 4.68E-01 | 7.52E-01 | 54 | 3.25E-01 | 2.85E-01 | 5.19E-01 | 4.81 | 3.77 | 8.15E-05 | -2.4 | 2.4 | -7.2 |
| | | 6.22E-01 | 7.52E-01 | 26 | 5.01E-01 | 5.62E-01 | 6.38E-01 | 4.22 | 3.44 | 2.86E-04 | -0.6 | 3.0 | -8.2 |
| | | 8.20E-01 | 8.82E-01 | 2 | 8.82E-01 | 7.96E-01 | 8.18E-01 | 3.78 | 3.18 | 7.40E-04 | 1.2 | 2.6 | -7.8 |
| **Hypometabolism - Control vs Repeated Cooling** | | | | | | | | | | | | | |
| 3.08E-01 | 3 | 4.50E-01 | 9.22E-01 | 58 | 3.07E-01 | 4.64E-01 | 9.28E-01 | 4.41 | 3.55 | 1.92E-04 | 3.8 | 1.6 | -2.2 |
| | | 7.24E-01 | 9.24E-01 | 12 | 6.62E-01 | 7.36E-01 | 9.28E-01 | 3.91 | 3.25 | 5.69E-04 | 1.2 | 0.8 | 2.4 |
| | | 8.34E-01 | 9.24E-01 | 1 | 9.24E-01 | 8.35E-01 | 9.28E-01 | 3.70 | 3.12 | 8.93E-04 | 0.6 | 0.0 | 3.0 |
| **Hypermetabolism - Control vs Repeated Restraint** | | | | | | | | | | | | | |
| 3.08E-01 | 3 | 5.14E-03 | 7.95E-03 | 654 | 2.65E-03 | 6.22E-02 | 1.12E-01 | 5.89 | 4.29 | 8.88E-06 | 0.8 | 4.4 | 3.2 |
| | | 7.97E-01 | 9.24E-01 | 4 | 8.20E-01 | 7.14E-01 | 8.06E-01 | 3.95 | 3.28 | 5.21E-04 | 0.2 | 6.4 | -2.4 |
| | | 8.34E-01 | 9.24E-01 | 1 | 9.24E-01 | 8.49E-01 | 9.73E-01 | 3.66 | 3.10 | 9.59E-04 | -3.4 | 5.8 | -4.8 |
| **Hypometabolism - Control vs Repeated Restraint** | | | | | | | | | | | | | |
| 3.08E-01 | 3 | 3.81E-01 | 4.87E-01 | 75 | 2.46E-01 | 6.53E-02 | 1.39E-01 | 5.86 | 4.28 | 9.49E-06 | 3.8 | 2.0 | -2.2 |
| | | 4.68E-01 | 4.87E-01 | 54 | 3.25E-01 | 5.09E-01 | 7.33E-01 | 4.32 | 3.50 | 2.32E-04 | 1.6 | 1.6 | -6.0 |
| | | 7.97E-01 | 8.20E-01 | 4 | 8.20E-01 | 8.03E-01 | 8.78E-01 | 3.77 | 3.17 | 7.63E-04 | 1.2 | 2.6 | -0.4 |
| **Hypermetabolism - Repeated Cooling vs Repeated Restraint** | | | | | | | | | | | | | |
| 1.93E-04 | 9 | 6.70E-01 | 9.24E-01 | 19 | 5.71E-01 | 4.05E-01 | 9.93E-01 | 4.53 | 3.62 | 1.49E-04 | -3.6 | 0.2 | -1.4 |
| | | 4.36E-01 | 9.24E-01 | 61 | 2.95E-01 | 4.76E-01 | 9.93E-01 | 4.39 | 3.54 | 2.02E-04 | 0.8 | 4.2 | 3.4 |
| | | 7.40E-01 | 9.24E-01 | 10 | 6.94E-01 | 6.37E-01 | 9.93E-01 | 4.09 | 3.36 | 3.83E-04 | 1.2 | 0.2 | 4.8 |
| | | 6.29E-01 | 9.24E-01 | 25 | 5.10E-01 | 7.12E-01 | 9.93E-01 | 3.95 | 3.28 | 5.16E-04 | -1.8 | 0.2 | 3.6 |
| | | 7.40E-01 | 9.24E-01 | 10 | 6.94E-01 | 7.56E-01 | 9.93E-01 | 3.87 | 3.23 | 6.20E-04 | 0.4 | 0.2 | 3.0 |
| | | 7.40E-01 | 9.24E-01 | 10 | 6.94E-01 | 7.76E-01 | 9.93E-01 | 3.83 | 3.20 | 6.76E-04 | 1.4 | -0.2 | 2.0 |
| | | 8.34E-01 | 9.24E-01 | 1 | 9.24E-01 | 8.51E-01 | 9.93E-01 | 3.66 | 3.10 | 9.70E-04 | -1.8 | 1.0 | 3.6 |
| | | 8.34E-01 | 9.24E-01 | 1 | 9.24E-01 | 8.53E-01 | 9.93E-01 | 3.65 | 3.10 | 9.82E-04 | -0.4 | 5.2 | -0.6 |
| | | 8.34E-01 | 9.24E-01 | 1 | 9.24E-01 | 8.55E-01 | 9.93E-01 | 3.65 | 3.09 | 9.89E-04 | 0.4 | 5.0 | 1.8 |
| **Hypometabolism - Repeated Cooling vs Repeated Restraint** | | | | | | | | | | | | | |
| 1.33E-01 | 4 | 4.08E-02 | 8.57E-02 | 341 | 2.14E-02 | 2.24E-01 | 4.68E-01 | 5.00 | 3.87 | 5.54E-05 | 1.2 | 2.6 | -7.8 |
| | | 4.82E-01 | 6.77E-01 | 51 | 3.39E-01 | 2.64E-01 | 4.68E-01 | 4.87 | 3.80 | 7.19E-05 | -2.4 | 2.4 | -7.2 |
| | | 8.20E-01 | 9.24E-01 | 2 | 8.82E-01 | 7.72E-01 | 9.13E-01 | 3.83 | 3.21 | 6.64E-04 | 3.8 | 2.2 | -2.2 |
| | | 8.34E-01 | 9.24E-01 | 1 | 9.24E-01 | 8.46E-01 | 9.63E-01 | 3.67 | 3.11 | 9.45E-04 | 1.2 | 4.0 | -8.0 |

**B**

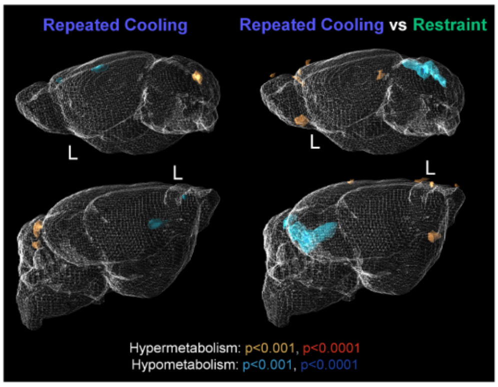


**Supplementary Figure 6: Voxel-wise brain analysis of repeated acute stress models**

**A**. Summarized quantitative results of voxel-wise analysis performed with statistical parametric mapping (SPM12) using a one-way ANOVA model with post-hoc t-contrasts. Corresponding images are shown in Fig. 2H and Supplementary Fig. 6B.

**B**. Voxel-wise exploratory comparison of normalized brain [$^{18}$F]FDG uptake showing differences between control (n = 8) and repeated cooling (n = 7) mice on the left and between repeated cooling (n = 7) and repeated restraint (n = 5) on the right, with thresholds of p < 0.001 and p < 0.0001 (uncorr.) to render significantly changed areas with the PMOD 3D tool (Ver. 4.4). The yellow and red areas indicate hypermetabolism, while the blue and dark blue areas indicate hypometabolism.

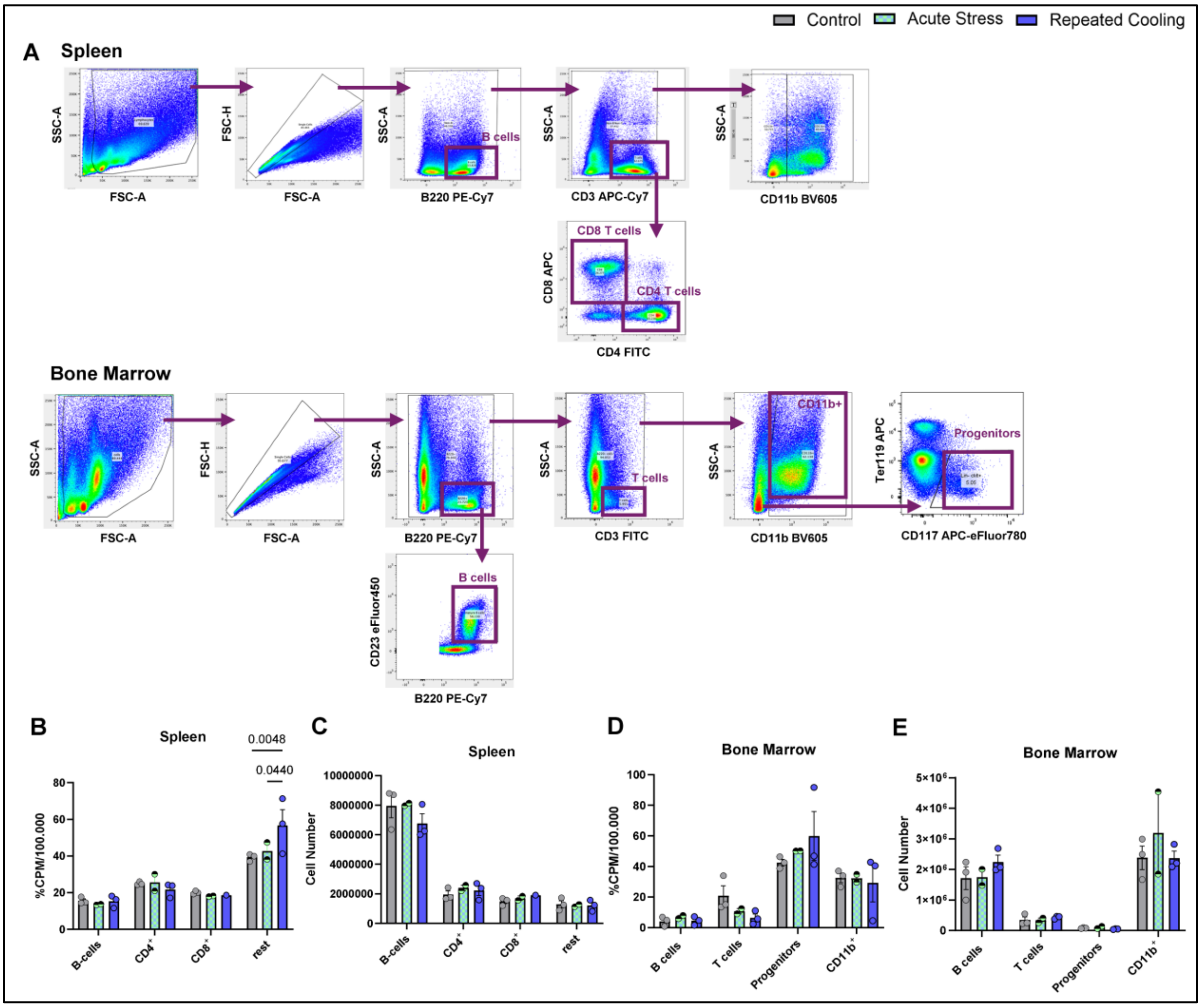


**Supplementary Figure 7: RadioFlow experiments reveal changes of the immune system in spleen and bone marrow after repeated cooling.**

**A**. Illustration of the gating strategy of spleen and bone marrow tissue used for RadioFlow experiments.

**B-E**. [$^{18}$F]FDG RadioFlow results of spleen (**B, C**) and bone marrow (**D, E**) as percentage of total counts per minute (%cpm) per 100,000 cells (**B, D**) and total cell number (**C, E**) in control (n = 3), acute stress (n = 2; cooling and restraint), and repeated cooling (n = 1-3) mice.

Data were analyzed using two-way ANOVA (**B-E**) with Tukey's post-hoc test. Data are presented as individual values and as mean ± SEM, with p-values indicated as numbers above the bars.

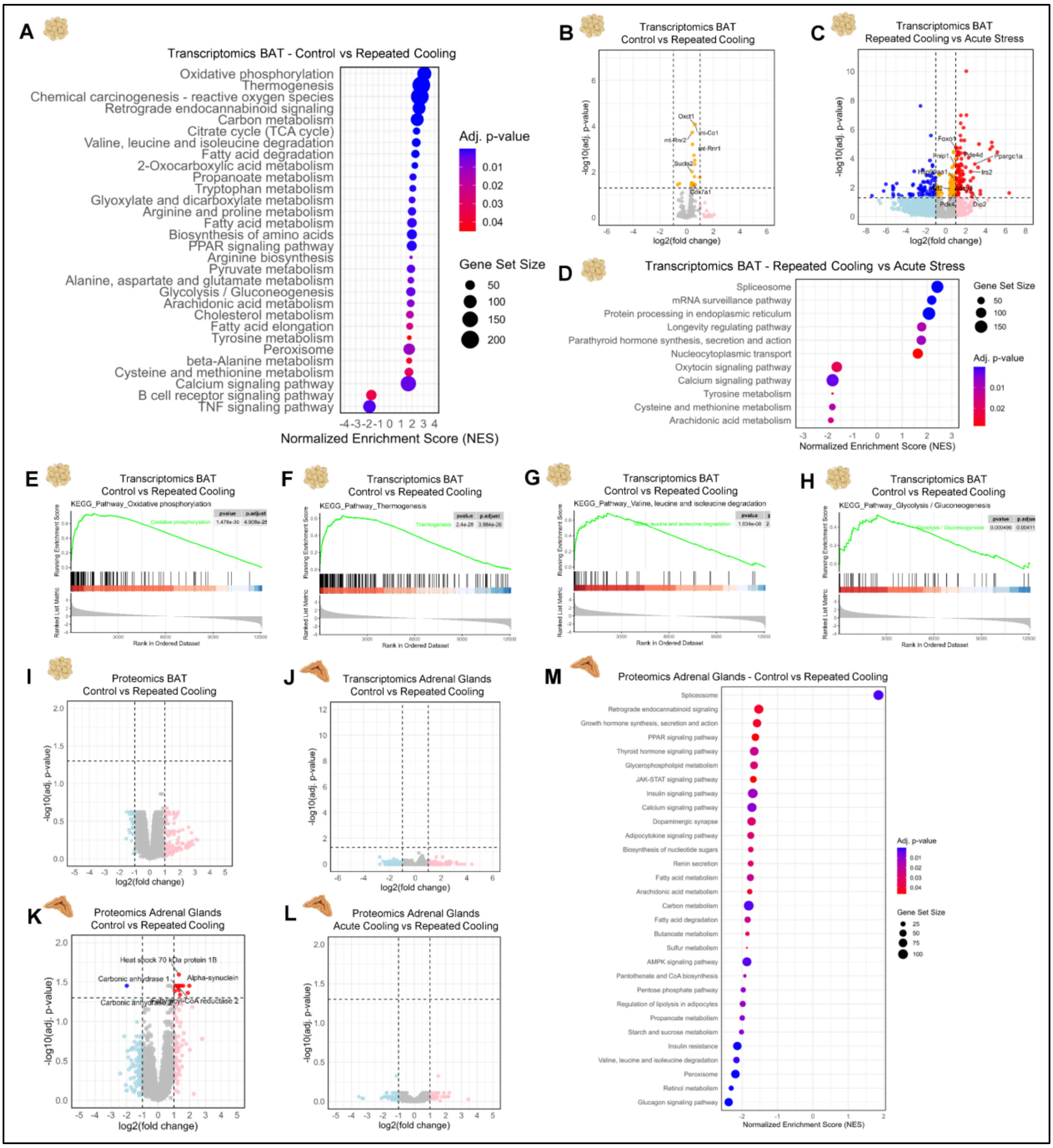


**Supplementary Figure 8: Bulk RNA-Seq and Proteomics analysis of Adrenal Glands and iBAT in repeated acute stress models**

**A**. KEGG pathway gene set enrichment analysis of bulk RNA-Sequencing results obtained via DESeq2 (**B**) of iBAT in control (n = 3) and repeated cooling (n = 5) mice illustrated as a bubble plot showing normalized enrichment score (NES), adjusted p-value (adj. p-value), and gene set size. The headings in the Omics datasets indicate the direction of the comparison (A vs. B), with A as the control condition.

**B, C**. Bulk RNA-sequencing results of iBAT in control (n = 3) vs repeated cooling (n = 5) (**B**) and repeated cooling (n = 5) vs acute stress (n = 3; cooling and restraint) (**C**) mice. Differential gene expression was analyzed using DESeq2 with two-sided Wald tests and Benjamini-Hochberg correction and illustrated as a volcano plot (downregulated: blue, downregulated (ns): light blue, upregulated: red, upregulated (ns): light red, |log2FC| ≤ 1 & p-adj. ≤ 0.05: orange, no change: grey).

**D**. KEGG pathway gene set enrichment analysis of bulk RNA-Sequencing results of iBAT (**C**) illustrated as a bubble plot showing normalized enrichment score (NES), adjusted p-value (adj. p-value), and gene set size.

**E-H**. Enrichment plots of Oxidative phosphorylation (**E**), Thermogenesis (**F**), Valine, Leucine, and Isoleucine degradation (**G**), and Glycolysis/Gluconeogenesis (**H**) KEGG pathways in control (n = 3) vs repeated cooling (n = 5) mice of iBAT via bulk RNA-Seq analysis.

**I**. Proteomics results of iBAT in control (n = 3) and repeated cooling (n = 6) mice, analyzed using LIMMA with two-sided moderated t-tests and Benjamini-Hochberg correction, illustrated as a volcano plot (legend see **B, C**).

**J**. Bulk RNA-sequencing results of adrenal glands in control (n = 5) vs repeated cooling (n = 5) mice. Differential gene expression was analyzed using DESeq2 with two-sided Wald tests and Benjamini-Hochberg correction and illustrated as a volcano plot (legend see **B, C**).

**K, L**. Proteomics results of adrenal gland tissue in control (n = 5) vs repeated cooling (n = 5) (**K**) and acute cooling (n = 3) vs repeated cooling (n = 5) (**L**) mice, analyzed using LIMMA with two-sided moderated t-tests and Benjamini-Hochberg correction and illustrated as a volcano plot (legend see **B, C**).

**M**. KEGG pathway gene set enrichment analysis of proteomics results of adrenal gland tissue (**K**) illustrated as a bubble plot showing normalized enrichment score (NES), adjusted p-value (adj. p-value), and gene set size.

Pictograms of organs in A-M were taken from BioRender. Vraka, C. (2026) https://BioRender.com/d1uqx40.

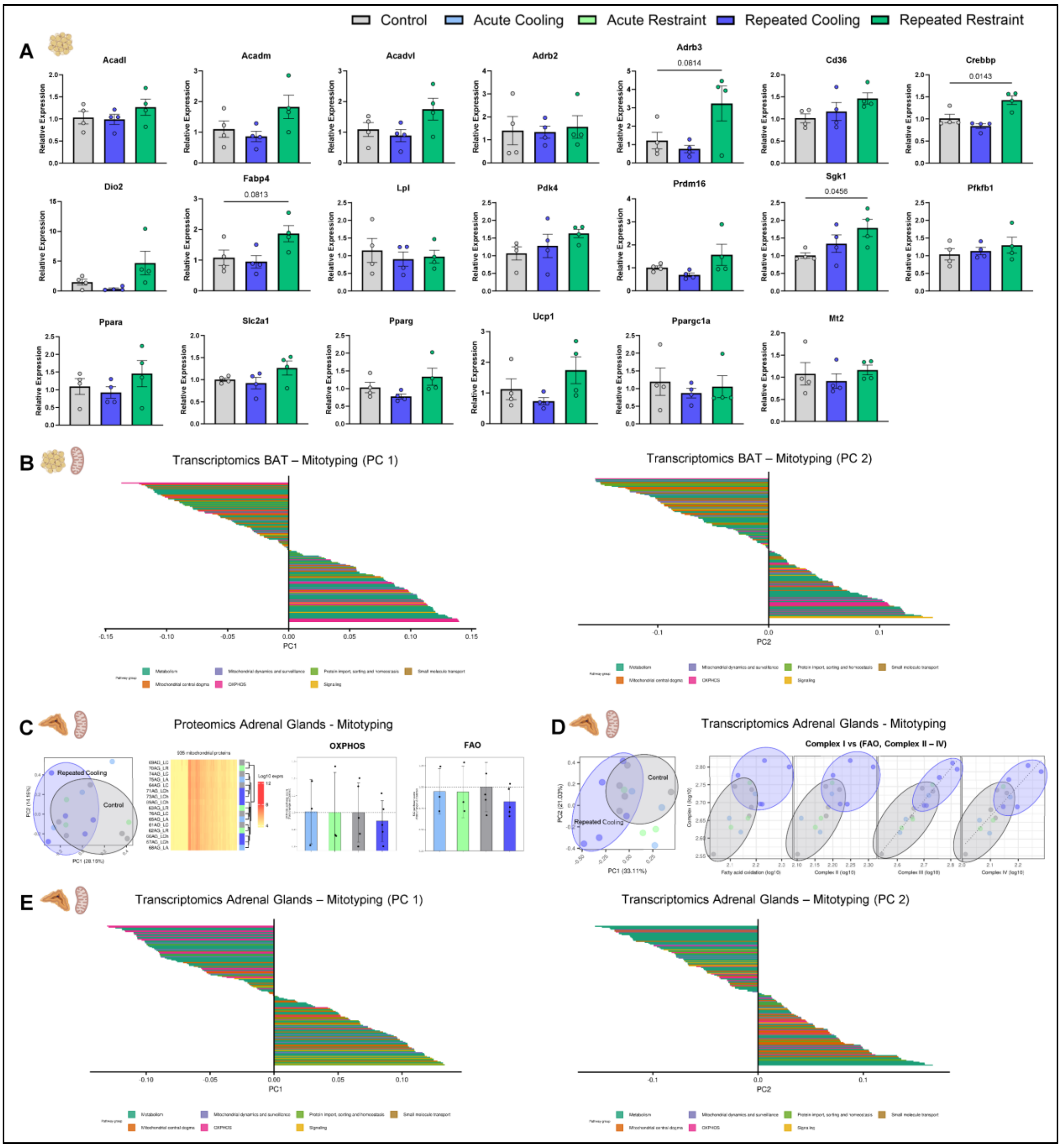


**Supplementary Figure 9: qPCR results of iBAT in repeated acute stress models and Mitotyping analysis of iBAT and adrenal glands**

**A**. Relative gene expression of iBAT tissue assessed via RT-qPCR in control (n = 4), repeated cooling (n = 4), and repeated restraint (n = 4) mice quantified via the ΔΔCt method.

**B**. Bars represent PCA loadings of individual MitoPathways based on mitochondrial Pathway Priority Scores (mitoPPS) derived from Transcriptomics data of iBAT in control (n = 3), acute stress (n = 3; cooling and restraint), and repeated cooling (n = 5) mice and are colored according to their MitoCarta Level 1 pathway classification. Larger absolute loading values indicate a stronger contribution of the respective pathway to the corresponding principal component. PC1 loadings from these subset-specific analyses indicate the pathways contributing most strongly to the dominant axis of mitochondrial variation within each subset.

**C**. Mitotyping analysis of Proteomics results in adrenal glands of control (n = 5), acute cooling (n = 3), acute restraint (n = 3), and repeated cooling (n = 5) mice. Left to right: 1. PCA analysis on mitochondrial Pathway Priority Scores (mitoPPS) displaying PC1–PC2 projections visualizing the major axes of variation in mitochondrial pathway prioritization. 2. Hierarchical clustering of samples according to 862 mitochondrial proteins. 3, 4. Oxidative phosphorylation (OXPHOS, 3.) and fatty acid oxidation (FAO, 4.) mitochondrial pathway scores per experimental group.

**D**. Mitotyping analysis of Transcriptomics results in adrenal glands of control (n = 5), acute cooling (n = 3), acute restraint (n = 3), and repeated cooling (n = 5) mice. Left to right: 1. PCA analysis on mitochondrial Pathway Priority Scores (mitoPPS) displaying PC1–PC2 projections visualizing the major axes of variation in mitochondrial pathway prioritization. 2. Plots showing OXPHOS (Complex I-IV) and FAO expression profiles of samples visualizing the molecular specialization of mitochondria.

**E**. PCA loadings as shown in (**B**) from Transcriptomics data of adrenal glands displayed in (**D**).

Data were analyzed using one-way ANOVA (**A**) with Dunnett's post-hoc test. Data are presented as individual values and as mean ± SEM, with p-values indicated as numbers above the bars. For further methodological details, refer to the Methods section.

Pictograms of organs in A-E were taken from BioRender. Vraka, C. (2026) https://BioRender.com/d1uqx40.

Control   Anesthesia d1   Surgery d1   Surgery d9

**A**

**Hypermetabolism - Control vs Anesthesia d1**

| set p | set c | cluster p(FWE-corr) | cluster p(FDR-corr) | cluster equivk | cluster p(unc) | peak p(FWE-corr) | peak p(FDR-corr) | peak T | peak equivZ | peak p(unc) | x,y,z {mm} | x,y,z {mm} | x,y,z {mm} |
|---|---|---|---|---|---|---|---|---|---|---|---|---|---|
| 3.56E-02 | 8 | 0.00E+00 | 1.34E-31 | 13815 | 1.67E-32 | 2.75E-07 | 1.44E-05 | 13.69 | 6.87 | 3.10E-12 | -0.8 | 6.4 | -2.4 |
| | | | | | | 9.34E-03 | 1.12E-02 | 6.93 | 4.95 | 3.80E-07 | 3.0 | 1.2 | 2.4 |
| | | 2.46E-02 | 2.69E-02 | 247 | 6.72E-03 | 1.14E-01 | 8.36E-02 | 5.49 | 4.28 | 9.55E-06 | -2.0 | 0.0 | 2.0 |
| | | 4.38E-01 | 3.11E-01 | 56 | 1.56E-01 | 6.79E-01 | 4.13E-01 | 4.24 | 3.56 | 1.85E-04 | 2.0 | 7.2 | -7.4 |
| | | 8.62E-01 | 7.11E-01 | 11 | 5.33E-01 | 7.68E-01 | 5.17E-01 | 4.09 | 3.47 | 2.62E-04 | 0.4 | 1.0 | 6.2 |
| | | 3.26E-01 | 2.84E-01 | 74 | 1.06E-01 | 8.06E-01 | 5.58E-01 | 4.02 | 3.42 | 3.08E-04 | 0.6 | 7.8 | -9.2 |
| | | 6.04E-01 | 4.00E-01 | 36 | 2.50E-01 | 9.26E-01 | 7.72E-01 | 3.75 | 3.24 | 5.93E-04 | -2.8 | 1.4 | -7.4 |
| | | 9.62E-01 | 8.81E-01 | 1 | 8.81E-01 | 9.67E-01 | 9.68E-01 | 3.58 | 3.13 | 8.80E-04 | 2.6 | 6.8 | -5.6 |
| | | 9.62E-01 | 8.81E-01 | 1 | 8.81E-01 | 9.74E-01 | 9.90E-01 | 3.54 | 3.10 | 9.70E-04 | -2.0 | -0.2 | -4.2 |

**Hypometabolism - Control vs Anesthesia d1**

| set p | set c | cluster p(FWE-corr) | cluster p(FDR-corr) | cluster equivk | cluster p(unc) | peak p(FWE-corr) | peak p(FDR-corr) | peak T | peak equivZ | peak p(unc) | x,y,z {mm} | x,y,z {mm} | x,y,z {mm} |
|---|---|---|---|---|---|---|---|---|---|---|---|---|---|
| 5.07E-01 | 4 | 0.00E+00 | 8.08E-36 | 16447 | 2.02E-36 | 6.00E-04 | 6.75E-03 | 8.58 | 5.56 | 1.32E-08 | 2.4 | 2.0 | -3.2 |
| | | 6.63E-01 | 5.86E-01 | 30 | 2.93E-01 | 7.00E-01 | 3.72E-01 | 4.20 | 3.54 | 2.00E-04 | 0.2 | 6.0 | -7.6 |
| | | 8.41E-01 | 6.60E-01 | 13 | 4.95E-01 | 8.73E-01 | 6.04E-01 | 3.89 | 3.34 | 4.26E-04 | -0.4 | 3.6 | 1.8 |
| | | 9.62E-01 | 8.81E-01 | 1 | 8.81E-01 | 9.74E-01 | 9.85E-01 | 3.54 | 3.10 | 9.77E-04 | -0.2 | 6.0 | -6.4 |

**Hypermetabolism - Control vs Surgery d1**

| set p | set c | cluster p(FWE-corr) | cluster p(FDR-corr) | cluster equivk | cluster p(unc) | peak p(FWE-corr) | peak p(FDR-corr) | peak T | peak equivZ | peak p(unc) | x,y,z {mm} | x,y,z {mm} | x,y,z {mm} |
|---|---|---|---|---|---|---|---|---|---|---|---|---|---|
| 1.71E-01 | 6 | 1.11E-15 | 1.74E-15 | 4723 | 2.91E-16 | 1.64E-05 | 1.15E-04 | 10.98 | 6.27 | 1.85E-10 | -0.2 | 6.2 | -2.4 |
| | | 1.92E-11 | 1.55E-11 | 2924 | 5.18E-12 | 7.26E-04 | 2.62E-03 | 8.46 | 5.52 | 1.66E-08 | -2.6 | 2.0 | 3.4 |
| | | 7.02E-10 | 3.79E-10 | 2338 | 1.89E-10 | 1.50E-03 | 2.93E-03 | 8.02 | 5.37 | 4.00E-08 | 1.0 | 4.0 | 4.0 |
| | | 5.50E-01 | 3.23E-01 | 42 | 2.15E-01 | 6.04E-01 | 2.90E-01 | 4.35 | 3.63 | 1.40E-04 | 0.0 | 8.0 | -9.8 |
| | | 7.03E-01 | 3.93E-01 | 26 | 3.28E-01 | 7.95E-01 | 4.55E-01 | 4.04 | 3.44 | 2.93E-04 | 0.8 | 0.0 | 2.8 |
| | | 8.92E-01 | 6.01E-01 | 8 | 6.01E-01 | 8.84E-01 | 5.99E-01 | 3.86 | 3.32 | 4.53E-04 | 1.2 | -0.2 | 1.6 |

**Hypometabolism - Control vs Surgery d1**

| set p | set c | cluster p(FWE-corr) | cluster p(FDR-corr) | cluster equivk | cluster p(unc) | peak p(FWE-corr) | peak p(FDR-corr) | peak T | peak equivZ | peak p(unc) | x,y,z {mm} | x,y,z {mm} | x,y,z {mm} |
|---|---|---|---|---|---|---|---|---|---|---|---|---|---|
| 7.16E-01 | 3 | 0.00E+00 | 8.99E-25 | 9366 | 3.00E-25 | 3.15E-03 | 2.31E-02 | 7.57 | 5.20 | 9.91E-08 | -3.2 | 3.0 | -4.0 |
| | | 5.95E-01 | 3.66E-01 | 37 | 2.44E-01 | 6.34E-01 | 3.06E-01 | 4.31 | 3.61 | 1.56E-04 | 0.8 | 4.4 | -7.6 |
| | | 9.13E-01 | 6.57E-01 | 6 | 6.57E-01 | 9.01E-01 | 6.24E-01 | 3.82 | 3.29 | 5.01E-04 | -1.4 | 4.0 | 0.0 |

**Hypermetabolism - Anesthesia vs Surgery d1**

| set p | set c | cluster p(FWE-corr) | cluster p(FDR-corr) | cluster equivk | cluster p(unc) | peak p(FWE-corr) | peak p(FDR-corr) | peak T | peak equivZ | peak p(unc) | x,y,z {mm} | x,y,z {mm} | x,y,z {mm} |
|---|---|---|---|---|---|---|---|---|---|---|---|---|---|
| 4.91E-03 | 10 | 3.47E-07 | 9.37E-07 | 1437 | 9.37E-08 | 3.00E-03 | 2.19E-02 | 7.60 | 5.21 | 9.36E-08 | 0.0 | 6.2 | -5.8 |
| | | 1.91E-01 | 1.43E-01 | 107 | 5.70E-02 | 1.48E-01 | 2.26E-01 | 5.34 | 4.20 | 1.36E-05 | -2.6 | 5.8 | -4.2 |
| | | 2.20E-03 | 2.97E-03 | 447 | 5.93E-04 | 1.70E-01 | 2.26E-01 | 5.25 | 4.15 | 1.66E-05 | -1.8 | 2.0 | 4.2 |
| | | 1.79E-01 | 1.43E-01 | 111 | 5.31E-02 | 5.22E-01 | 6.12E-01 | 4.48 | 3.71 | 1.03E-04 | -0.8 | 5.4 | -1.4 |
| | | 8.30E-01 | 8.81E-01 | 14 | 4.78E-01 | 8.76E-01 | 7.82E-01 | 3.88 | 3.33 | 4.32E-04 | 0.6 | 4.0 | 4.2 |
| | | 9.13E-01 | 8.81E-01 | 6 | 6.57E-01 | 9.05E-01 | 7.82E-01 | 3.81 | 3.28 | 5.15E-04 | 2.0 | 6.2 | -3.0 |
| | | 9.42E-01 | 8.81E-01 | 3 | 7.68E-01 | 9.25E-01 | 7.82E-01 | 3.75 | 3.24 | 5.88E-04 | -0.6 | 3.8 | 3.0 |
| | | 9.62E-01 | 8.81E-01 | 1 | 8.81E-01 | 9.26E-01 | 7.82E-01 | 3.75 | 3.24 | 5.96E-04 | 1.0 | 3.4 | 5.0 |
| | | 9.62E-01 | 8.81E-01 | 1 | 8.81E-01 | 9.66E-01 | 9.21E-01 | 3.59 | 3.13 | 8.68E-04 | 0.8 | 4.2 | 3.6 |
| | | 9.62E-01 | 8.81E-01 | 1 | 8.81E-01 | 9.67E-01 | 9.21E-01 | 3.58 | 3.13 | 8.84E-04 | 0.0 | 1.2 | 0.6 |

**Hypometabolism - Anesthesia vs Surgery d1**

| set p | set c | cluster p(FWE-corr) | cluster p(FDR-corr) | cluster equivk | cluster p(unc) | peak p(FWE-corr) | peak p(FDR-corr) | peak T | peak equivZ | peak p(unc) | x,y,z {mm} | x,y,z {mm} | x,y,z {mm} |
|---|---|---|---|---|---|---|---|---|---|---|---|---|---|
| 1.71E-01 | 6 | 8.33E-07 | 1.35E-06 | 1322 | 2.25E-07 | 1.95E-02 | 4.90E-02 | 6.50 | 4.76 | 9.58E-07 | -5.6 | 2.6 | -2.0 |
| | | 1.16E-04 | 9.42E-05 | 737 | 3.14E-05 | 1.07E-01 | 1.73E-01 | 5.53 | 4.30 | 8.70E-06 | 4.4 | 3.2 | -3.4 |
| | | 1.10E-01 | 6.29E-02 | 142 | 3.15E-02 | 4.23E-01 | 3.55E-01 | 4.65 | 3.81 | 6.90E-05 | -2.4 | 4.0 | -7.6 |
| | | 8.92E-01 | 8.81E-01 | 8 | 6.01E-01 | 9.38E-01 | 8.49E-01 | 3.71 | 3.21 | 6.53E-04 | -4.2 | 1.6 | -3.6 |
| | | 9.52E-01 | 8.81E-01 | 2 | 8.17E-01 | 9.53E-01 | 8.79E-01 | 3.65 | 3.17 | 7.55E-04 | 3.6 | 1.6 | 0.4 |
| | | 9.62E-01 | 8.81E-01 | 1 | 8.81E-01 | 9.72E-01 | 9.60E-01 | 3.55 | 3.11 | 9.41E-04 | -4.4 | 0.8 | -0.2 |

**B**

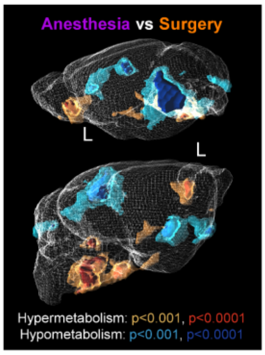


**Supplementary Figure 10: Voxel-wise brain analysis in surgery models**

**A**. Summarized quantitative results of voxel-wise analysis performed with statistical parametric mapping (SPM12) using a one-way ANOVA model with post-hoc t-contrasts. Corresponding images are shown in Fig. 3L and Supplementary Fig. 10B.

**B**. Voxel-wise exploratory comparison of normalized brain [$^{18}$F]FDG uptake performed with statistical parametric mapping (SPM12) using a one-way ANOVA model with post-hoc t-contrasts and thresholds of $p < 0.001$ and $p < 0.0001$ (uncorr.) to render significantly changed areas with the PMOD 3D tool (Ver. 4.4). Images show only the significant changes between anesthesia d1 ($n = 6$) and surgery d1 ($n = 6$) mice. The yellow and red areas indicate hypermetabolism, while the blue and dark blue areas indicate hypometabolism.

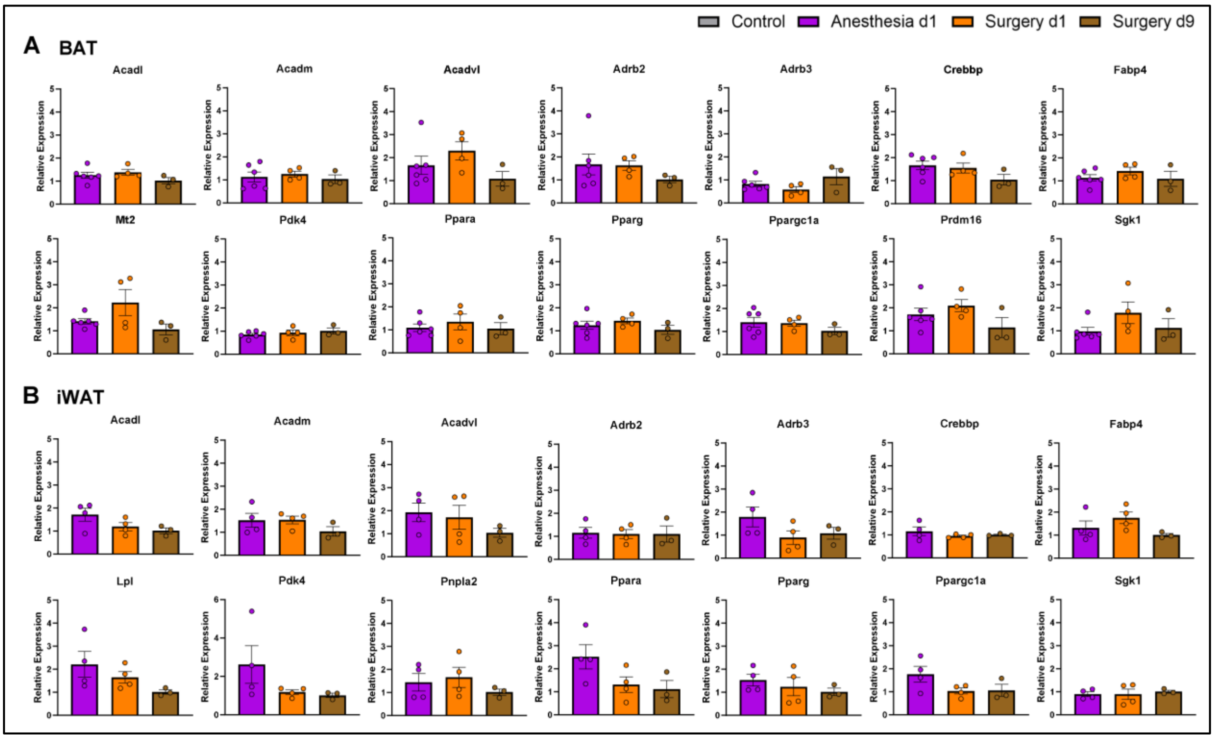


**Supplementary Figure 11: qPCR results of iBAT and iWAT in surgery models**

**A, B**. Relative gene expression of iBAT (**A**) and iWAT (**B**) assessed via RT-qPCR in anesthesia d1 (A, n = 6; B, n = 4), surgery d1 (n = 4), and surgery d9 (n = 3) mice, quantified via the ΔΔCt method.

Data were analyzed using one-way ANOVA (**A, B**) with Tukey's post-hoc test. Data are presented as individual values and as mean ± SEM, with p-values indicated as numbers above the bars.

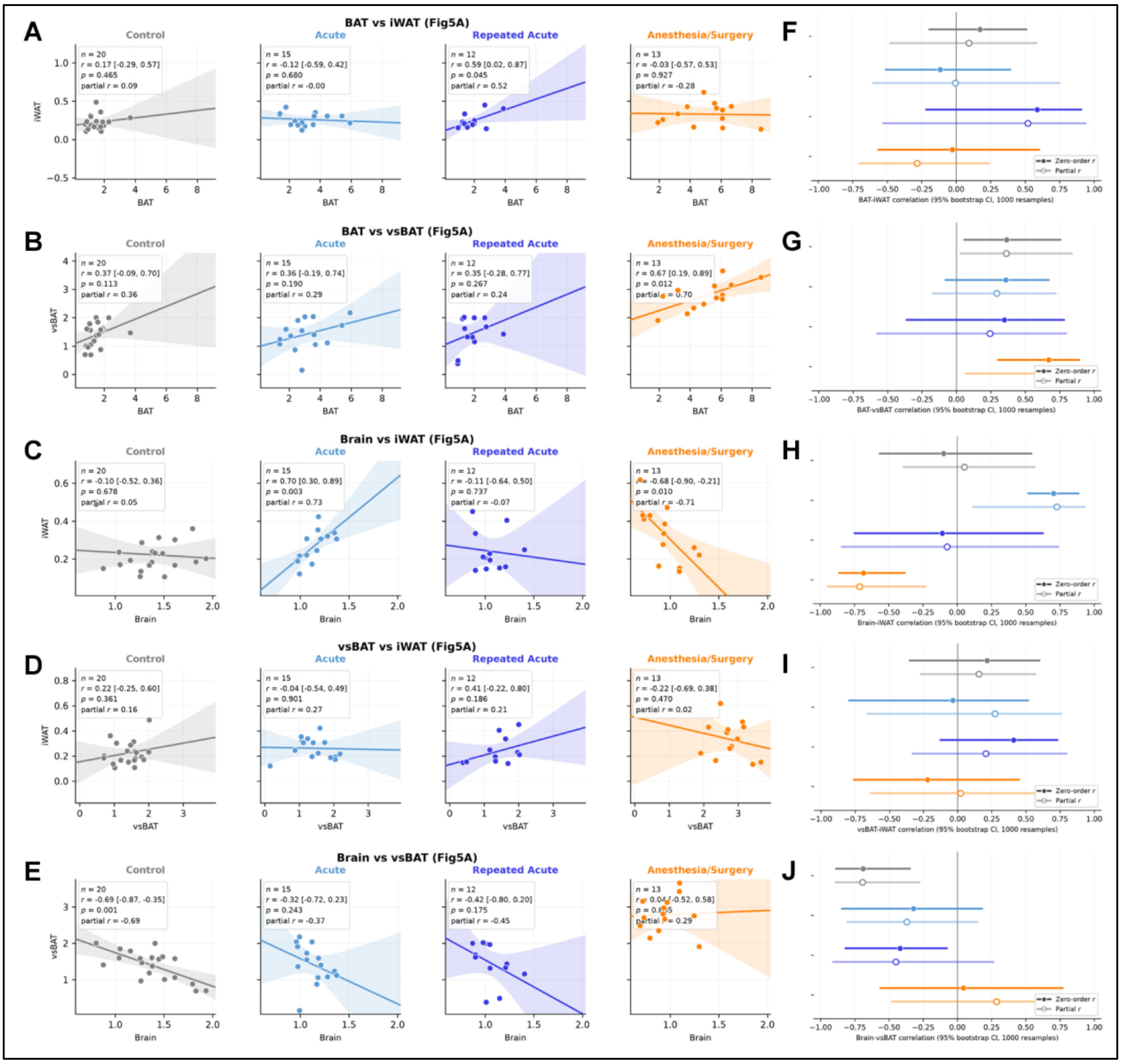


**Supplementary Figure 12: Results of Brain-Adipose Tissue Network Analysis**

**A-E**. Scatter plots illustrating the correlation between BAT - iWAT (A), BAT - vsBAT (B), Brain - iWAT (C), vsBAT - iWAT (D), and Brain - vsBAT (E) (all $SUV_{mean}$) for each experimental group from **Fig. 5A:** control (n = 20), acute (n = 15), repeated acute (n = 12), and anesthesia/surgery (n = 13). Shaded bands indicate 95% confidence intervals of the fitted regression lines.

**F-J**. Forest plots showing each group's zero-order correlation coefficient (filled symbols) and partial correlation coefficient (empty symbols), with their bootstrap estimates (95% confidence intervals, 1000 resamples), of BAT - iWAT (F), BAT - vsBAT (G), Brain - iWAT (H), vsBAT - iWAT (I), and Brain - vsBAT (J) from **Fig. 5A**.

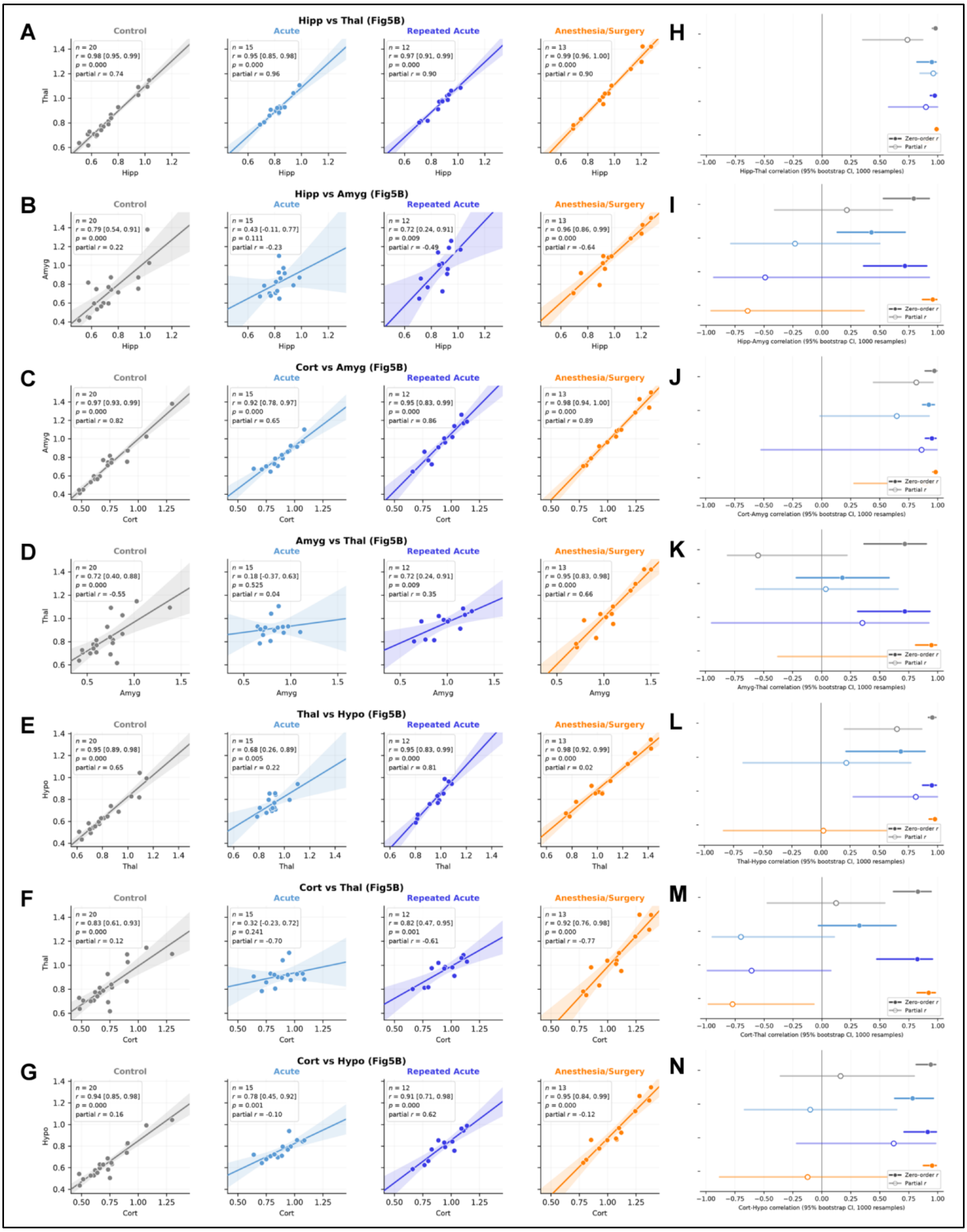


**Supplementary Figure 13: Results of Brain Region Network Analysis**

**A-G**. Scatter plots illustrating the correlation between Hipp – Thal (A), Hipp – Amyg (B), Cort – Amyg (C), Amyg – Thal (D), Thal – Hypo (E), Cort – Thal (F), Cort – Hypo (G) (all $SUV_{ratio}$) for each experimental group from **Fig. 5D:** control (n = 20), acute (n = 15), repeated acute (n = 12), and anesthesia/surgery (n = 13). Shaded bands indicate 95% confidence intervals of the fitted regression lines.

**H-N**. Forest plots showing each group's zero-order correlation coefficient (filled symbols) and partial correlation coefficient (empty symbols), with their bootstrap estimates (95% confidence intervals, 1000 resamples) of Hipp – Thal (H), Hipp – Amyg (I), Cort – Amyg (J), Amyg – Thal (K), Thal – Hypo (L), Cort – Thal (M), Cort – Hypo (N) from **Fig. 5D**.
